# Blood glucose homeostasis in patients with mild to moderate Spinocerebellar Ataxia type 2

**DOI:** 10.64898/2026.01.25.701567

**Authors:** Raúl Aguilera-Rodríguez, Dennis Almaguer-Gotay, Amarilis Álvarez-Sosa, Marilennis Anidos-Machado, Yanelis Silva-Ricardo, Dany Cuello-Almarales, Annelié Estupiñán-Rodríguez, Luis E. Almaguer-Mederos

**Author notes:** joint first authorship (Raúl Aguilera-Rodríguez, Dennis Almaguer-Gotay). Correspondence to Luis E. Almaguer-Mederos.

## Abstract

**Background:** Spinocerebellar ataxia type 2 (SCA2) is a neurodegenerative disorder that shows cerebellar glucose hypometabolism, systemic hypermetabolism and weight loss. This study aimed to explore novel molecular biomarker candidates for SCA2, based on the assessment of blood glucose homeostasis.

**Methods:** A case-control and correlational study was conducted in 79 Cuban patients with SCA2 and 83 sex- and age-matched control subjects during a fasting state. A subset of 20 SCA2 patients and 19 control individuals underwent an oral glucose tolerance test (OGTT). Several indices for assessing blood glucose homeostasis, derived from the fasting state or the OGTT, were included in the study.

**Results:** Fasting glucose levels showed a small increase, whereas the QUICKI index for insulin sensitivity was slightly decreased among patients. Markers of glucose homeostasis derived from the OGTT were no different between patients and controls. Fasting insulin levels, QUICKI, McAuley’s, HOMA2-%S, HOMA2-IR, and HOMA2-%β indices showed weak to moderate correlations with markers of body composition. McAuley’s, TyG, HOMA2-IR, and HOMA2-%β indices correlated with the age at onset, progression rate, or INAS count.

**Conclusions:** SCA2 patients with mild to moderate ataxia do not have any major alteration in blood glucose homeostasis. Markers of glucose homeostasis associate with body composition and has modifying effects on disease severity and progression rate in patients with SCA2, with McAuley’s index showing effects more consistently. Further studies are needed to describe the changes in blood glucose homeostasis in the different stages of disease, and to confirm the validity of McAuley’s index as a candidate biomarker for SCA2 clinical severity and progression.

## Introduction

Blood glucose homeostasis, a tightly regulated process of maintaining blood glucose at a steady-state level, is critical for properly preserving a healthy cellular energy metabolism. This is particularly relevant to the brain, given that brain cells, mostly neurons, rely on the uptake and processing of glucose as their primary energy source for their highly energy-demanding metabolic processes and physiological activities (1, 2). Beyond its significance as a chemical energy provider, glucose metabolism in the brain is also important for the synthesis of amino acids, methyl donors derived from one-carbon metabolism, glycolipids, glycoproteins, precursors of neurotransmitters and neuromodulators, oxidative stress management, and neuroplasticity (1, 3, 4). Not surprisingly, therefore, altered brain glucose metabolism was linked to neurodegenerative disorders including Alzheimeŕs disease (AD) (5), Parkinsońs disease (PD) (6), prion diseases (7), amyotrophic lateral sclerosis (8), and frontotemporal dementias (9). Furthermore, brain glucose dyshomeostasis was also reported for polyglutamine disorders (polyQ), including Huntingtońs disease (HD) (10–12), spinocerebellar ataxias (SCA) type 1, 2, 3, 6, 7, and 17 (13–16), and dentatorubral-pallidoluysian atrophy (17).

Further evidence supporting a pathologic role for glucose dyshomeostasis in polyQ diseases comes from the study of peripheral blood surrogate biomarkers in patients and animal models. Initial reports on an augmented incidence of diabetes mellitus (DM) and altered response to glucose challenge during the oral glucose tolerance test (OGTT) among HD patients (18, 19), were followed by proof of a diabetic phenotype in HD (R6/2) transgenic mice, resulting from an age-dependent reduction of insulin transcript expression (20). Besides, increased insulin resistance, reduced insulin sensitivity, and decreased early-phase insulin secretion capacity have been reported in normoglycemic HD and SCA1 patients, suggesting an increased risk of type 2 diabetes (21, 22). In addition, anecdotal findings of increased blood plasma glucose levels in HD, SCA1 and SCA3 patients (23–25), decreased serum insulin levels, and increased insulin sensitivity along with reduced insulin resistance in SCA3 patients (26), and a marked decrease in blood glucose levels in mid-stage disease SCA7 266Q knock-in mice (27) were reported. Furthermore, administration of the hypoglycemic agent exendin-4 improved insulin sensitivity and secretion, motor coordination, and extended the lifespan of N171-82Q transgenic mice, suggesting targeting of glucose metabolism as a therapeutic strategy for HD (28). Interestingly, increased serum insulin levels suggestive of insulin resistance were observed in an Ataxin-2 deficient mouse (Sca2^-/-^) (29), and it was proposed that genetic variants at the chromosomal locus of human *ATXN2* carry a risk for DM (30). However, no study has been conducted on blood glucose homeostasis and its clinical relevance in patients with SCA2.

Spinocerebellar ataxia type 2 (MIM: 183090) is an autosomal-dominant polyQ disorder that reaches the highest prevalence worldwide in Holguin province, Cuba. Its genetic etiology consists of a dynamic CAG repeat expansion mutation in the first exon of the *ATXN2* gene (cytogenetic band: 12q24.12) (31–33). *ATXN2* CAG repeat expansions beyond 31 repeats cause the typical SCA2 cerebellar syndrome, while expansions in the range of 27 to 33 repeats increase the disease risk or progression of amyotrophic lateral sclerosis (ALS) (34–36), frontotemporal dementia (FTD) (37, 38), idiopathic levodopa-responsive Parkinson’s disease (39), and progressive supranuclear palsy (PSP) (40).

SCA2 clinical presentation includes gait ataxia, dysmetria, dysdiadochokinesia, and dysarthria, frequently accompanied by decreased tendon reflexes, leg cramps, kinetic or postural tremor, abnormal eye movements with slowed saccades progressing to nuclear ophthalmoplegia, and reduced body mass index and survival (41–45). The highly variable clinical presentation depends heavily on the length of the CAG repeat tract, with longer repeats causing an earlier disease onset and a faster progression (46, 47). Heritability estimates for the age at onset suggested an important contribution of additional genetic factors to disease severity (48).

The *ATXN2* gene encodes Ataxin-2 protein, a cytoplasmic RNA-binding factor involved in the regulation of RNA metabolism, mRNA translation, endocytosis, cellular growth and stress responses, cytoskeletal dynamics, and the inhibition of the mechanistic target of rapamycin complex 1 (mTORC1) signaling (49–56). Upon the unstable expansion of its polyQ domain, the Ataxin-2 protein acquires a toxic gain of function, leading to multi-level brain atrophy by promoting protein aggregation, transcriptional dysregulation, calcium signaling dysregulation, autophagy dysfunction, and oxidative stress (44, 49, 57–62).

Although significant progress was achieved in understanding the mechanisms of disease, and promising results were obtained in pre-clinical trials based on antisense oligonucleotides (63, 64) or Cas13 CRISPR effectors (65) against *Atxn2* mRNA, the full elucidation of the molecular processes and events involved in SCA2 pathophysiology is critical for identifying reliable surrogate biomarkers and additional targets for therapy. In this study, we assessed blood glucose homeostasis in a cohort of normoglycemic patients with SCA2 and healthy control subjects, considering their relationship with molecular and clinical findings.

## Methods

### Study Design

A case-control and correlational study was conducted to assess markers of blood glucose homeostasis in 79 Cuban patients with SCA2 and 83 healthy control subjects during fasting. A subset of 20 SCA2 patients and 19 healthy controls underwent an OGTT. Healthy control subjects were selected from the same geographical and ethnic backgrounds as the patients with SCA2 and were matched for age (±2 years) and sex. Only normoglycemic individuals with fasting serum glucose levels between ≥3.9 and <5.6 mmol/L (66) and without a previous diagnosis of prediabetes or diabetes mellitus or intestinal malabsorption were included in the study. SCA2 patients who were confined to a wheelchair or bed (disease stage 3) (67) were not included in the study. The Ethics Committee of the Center for the Investigation and Rehabilitation of Hereditary Ataxias approved the study protocol, and a consent form was obtained from patients according to the Declaration of Helsinki.

### Clinical and Genetic Assessments

Diagnosis of SCA2 relied on family history and clinical and molecular criteria. Clinical diagnosis was based on the presence of gait ataxia, dysdiadochokinesis, dysarthria, dysmetria, dysphagia, and slow saccades. Information was also obtained regarding sex, age, and disease duration in years from disease onset to the latest examination. Age at onset was defined as the onset of motor impairment, and it was estimated through a retrospective review of patients’ clinical records and interviews with patients and close relatives. The Scale for the Assessment and Rating of Ataxia (SARA score) was used for assessing ataxia severity (68); this scale varies from zero to 40 points, increasing with ataxia severity. The progression rate of ataxia at presentation was defined as the ratio between the SARA score and disease duration. The INAS count was used for assessing the severity of non-ataxia signs (69). The disease stage was assessed as previously specified (67), and the CAG repeat length at the *ATXN2* gene was determined as reported (70).

### Body composition measurements

A trained researcher quantified weight (expressed in kilograms, kg) and height (expressed in meters, m) following standardized procedures using a mechanical High-Quality Dial Body Scale (Perlong Medical Equipment Co., Ltd., Nanjing, China). The body mass index (BMI) was defined as the ratio between body weight and the square of the body height, and it was expressed in units of kg/m^2^. The waist circumference (expressed in centimeters, cm) was measured using a flexible, non-stretchable tape measure, following a standardized protocol (71).

### Assessment of blood glucose homeostasis

Blood samples were drawn between 08:00 and 10:00 a.m., after overnight fasting, considering known temporal variations of glucose and insulin levels due to diurnal rhythm (72). This was followed by an OGTT to assess glucose tolerance status and OGTT-related insulin release. During the OGTT, blood samples were collected immediately before (at minute 0) and 30, 60, 90, and 120 minutes after ingestion of a 75g oral glucose load in 3–5 minutes, after an overnight fast. The serum was separated by centrifugation and stored at − 20 °C until use. Glucose and triglyceride serum levels were measured by spectrophotometry on a Hitachi 902 Automatic Analyzer (Roche, Germany) following the manufactureŕs instructions. Serum insulin levels were quantified using the Insulin [^125^I] immunoradiometric assay (IRMA) kit (Izotop #RK-400CT, Hungary). The inter-assay coefficients of variation were less than 5%.

Several indices for the assessment of blood glucose homeostasis, based on serum glucose, insulin, or triglyceride levels, were included in the study. Fasting insulin sensitivity was assessed by QUICKI (73), McAuley index (Mffm/I) (McAuley et al., 2001), and the homeostasis model assessment of sensitivity (HOMA2-%S) (74). Fasting insulin resistance was evaluated by the Triglyceride-Glucose index (TyG) (Simental-Mendía, Rodríguez-Morán, & Guerrero-Romero, 2008), and the homeostasis model assessment of insulin resistance (HOMA2-IR) (74). Besides, fasting β-cell function was assessed by the homeostasis model assessment of beta-cell function (HOMA2-%β) (Levy, Matthews, & Hermans, 1998). HOMA2 indices were determined using the HOMA Calculator (v2.2.3).

Well-established indices derived from the OGTT were also quantified. The Gutt insulin sensitivity index (ISI0, 120) (75), Matsuda insulin sensitivity index (MI) (76), Stumvoll insulin sensitivity index (Stumvoll−ISI) (77), and Stumvoll metabolic clearance rate (Stumvoll-MCR) (77) were included as surrogate markers of insulin sensitivity/resistance. Additionally, the insulinogenic index (IGI) (78), Stumvoll phase 1 (PH1) and 2 (PH2) (77), and the ratio of the area under the curve for insulin from 0 to 120 minutes to the area under the curve for glucose from 0 to 120 minutes (AUC-i/g) (76), were used as surrogate markers of insulin secretion. Besides, the product of the insulinogenic and Matsuda indices (IGI x MI), AUC-i/g and Matsuda indices (AUC-i/g x MI), and the ratio of the insulinogenic index and fasting insulin levels (IGI / FI), were included in the study as composite measures mathematically analogous to the disposition index, indicating the ability of β‐cells to compensate for insulin resistance (79–81).

### Statistical Analysis

Descriptive statistics were applied to the quantitative variables under study. The Kolmogorov-Smirnov test was used to test all variables for normal distribution. The χ^2^ test was used to make comparisons for qualitative variables, whereas the Student’s t-test was used to establish comparisons for continuous variables between patients with SCA2 and control subjects. In cases when the homoscedasticity assumption did not hold, the Welch’s t-test was used instead. For comparison purposes between patients with SCA2 and control subjects, the markers included in the study were corrected for age, sex, or BMI by linear regression. Glucose and insulin curves during OGTT were analyzed using a General Linear Model for repeated measures and multiple comparison tests. Correlation analyses were done using the Pearson correlation coefficient. Stepwise multiple linear regression analyses were performed in patients with SCA2 in the search for factors relevant to the age at disease onset, SARA score, progression rate, or INAS count, considering *ATXN2* expanded alleles, disease duration, and markers of blood glucose homeostasis as independent variables. All tests were two-tailed, and statistical significance was defined as p < 0.05. The Holm-Bonferroni method was used to correct for multiple hypothesis testing. The SPSS software version 22.0 (SPSS Inc., Chicago, IL, USA) was used for all statistical analyses.

## Results

In light of the evidence of glucose cerebellar hypometabolism in SCA2 patients (14), increased insulin serum levels in Ataxin-2 deficient mice (Sca2^-/-^) (29), and on the assumption that altered blood glucose homeostasis may occur in SCA2 patients as a consequence of polyQ-expanded Ataxin-2 partial loss-of-function, we explored several markers of blood glucose homeostasis from fasting state in SCA2 patients relative to healthy control individuals. Patients in stage 1 (mild ataxia, n=37) or 2 (moderate ataxia, n=42) were similarly represented in the sample. The distributions of sex, age, body weight, waist circumference, and BMI showed no significant differences between SCA2 patients and control individuals (Table 1).

**Table 1.**
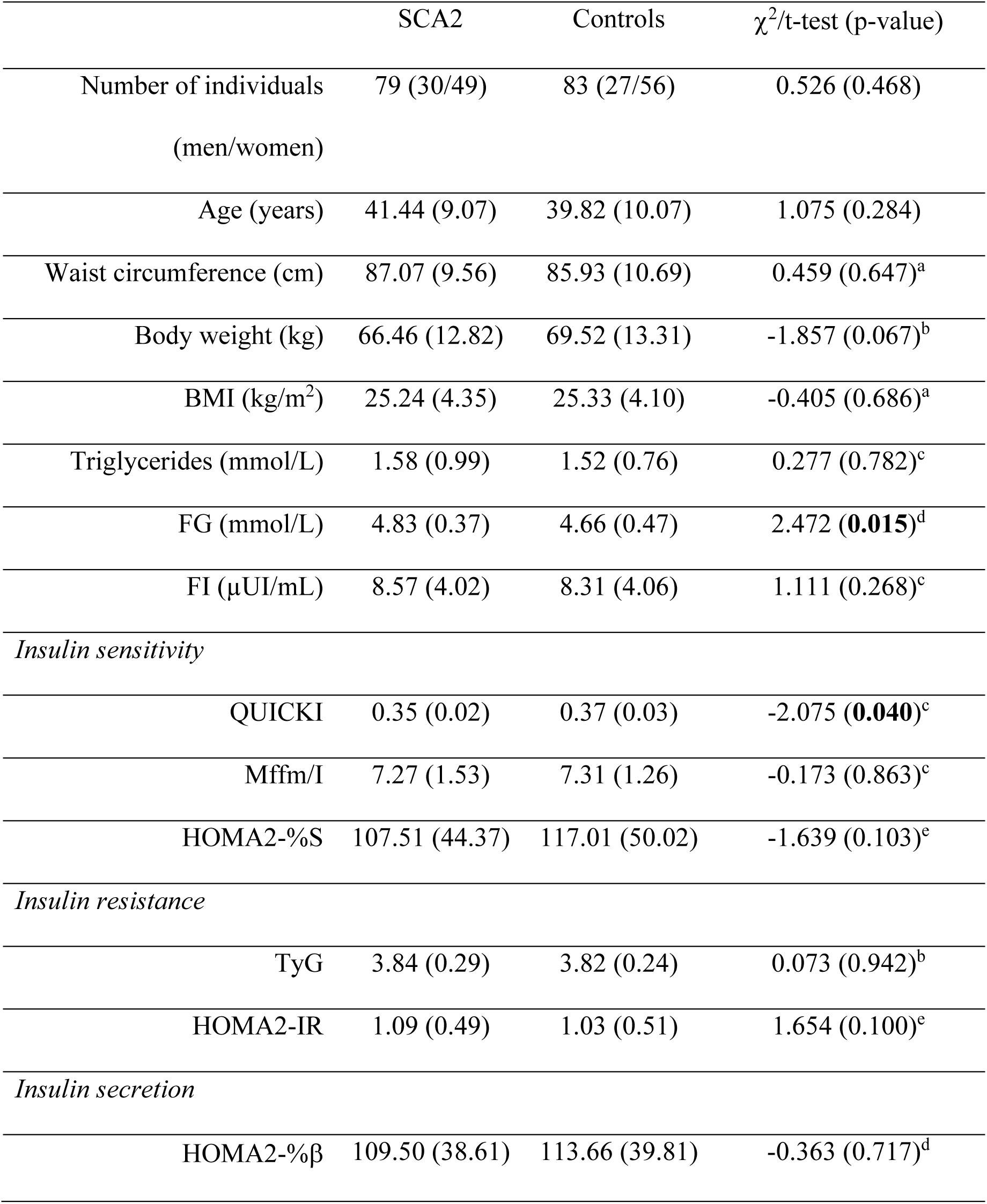
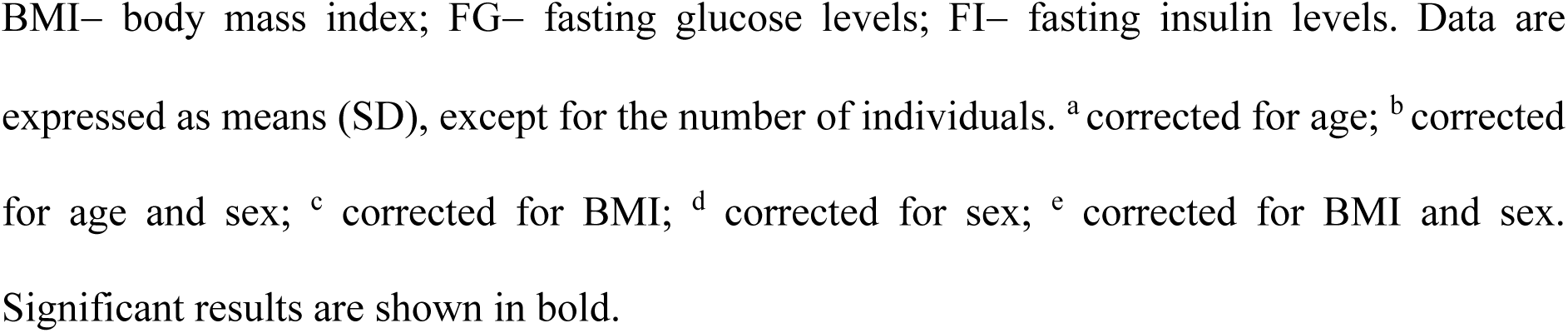
Descriptive statistics and mean comparisons for demographic variables and markers of glucose metabolism from the fasting state, stratified by clinical status.

### Blood glucose homeostasis is preserved in patients with SCA2

Neither fasting triglycerides nor insulin serum levels showed significant differences between patients and controls. Conversely, fasting glucose levels showed a small increase among patients after correction for sex; however, this difference did not remain significant after correction for multiple comparisons (corrected p-value = 0.210). Besides, patients were no different from controls for insulin secretion (based on the HOMA2-%β index), nor for insulin sensitivity/resistance (based on the TyG, Mffm/I, HOMA2-%S, and HOMA2-IR indices). Conversely, the QUICKI index for insulin sensitivity was slightly decreased with nominal significance among patients after correction for BMI, probably because of the increased glucose levels observed in the patient’s group (Table 1). Once again, this difference did not remain significant after correction for multiple comparisons (corrected p-value = 0.520).

A closer examination of the distribution of QUICKI index values across comparison groups, considering QUICKI values of <0.35 as indicative of possible insulin resistance (82, 83), produced no significant differences between patients and control individuals (χ^2^=0.960; p=0.327). Similar results were obtained when comparing additional insulin sensitivity, resistance, and secretion indices according to previously reported cut-off values (84–88) (Table S1).

As a further effort to uncover likely abnormalities in blood glucose homeostasis in SCA2 patients, we assessed the response of 20 SCA2 patients (15 females) and 19 control individuals (13 females) to a glucose load challenge during the OGTT. Comparison groups were no different from each other regarding sex (χ^2^=0.201; p=0.654), age (t=0.467; p=0.644), and BMI (t=0.302; p=0.765) distributions. Glucose levels during the OGTT were higher in the SCA2 patients relative to control individuals at every time point, but did not reach statistical significance (Figure 1A; Table S2). Conversely, insulin levels during the OGTT were lower in the SCA2 patients relative to control individuals at every time point after the oral glucose load, but no statistical significance was achieved (Figure 1B; Table S2).

**Figure 1.**
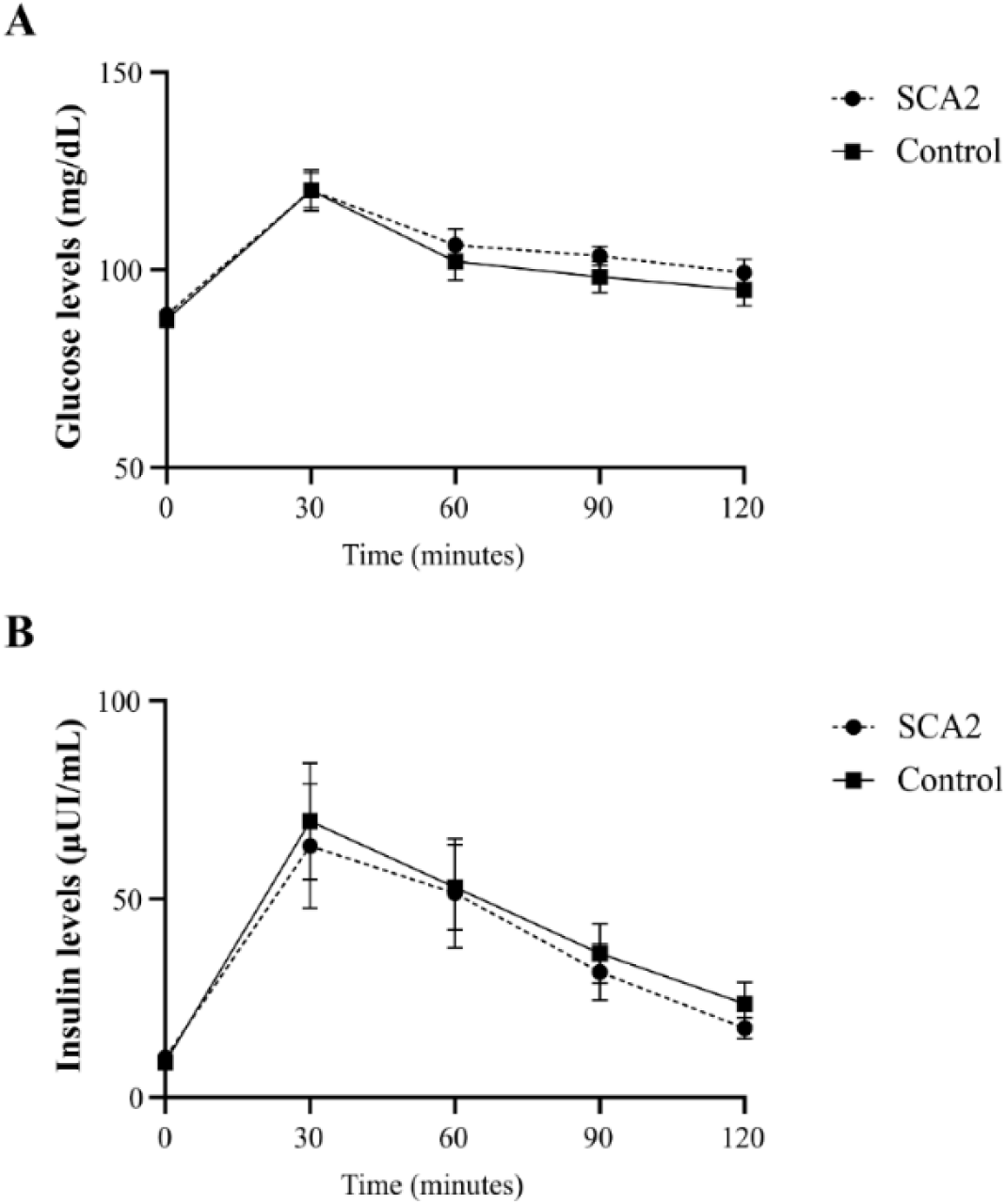
Oral glucose tolerance test (OGTT) in SCA2 patients and control individuals. **(A)** Mean serum glucose levels at different time points during OGTT. **(B)** Mean serum insulin levels at different time points during OGTT. Error bars represent SEM.

Next, we explored several indices of blood glucose homeostasis derived from the OGTT. All surrogate markers of insulin sensitivity were reduced in the patients relative to control individuals, but none of them reached statistical significance. Similarly, the insulinogenic and Stumvoll phase I (PH1) indices, as surrogate markers of early-phase insulin secretion, and the Stumvoll phase II (PH2) index, as an estimator of the sustained insulin secretion after the initial burst (77), were non-significantly reduced in the patients. The ratio between the area under the insulin curve and the area under the glucose curve (AUC i/g) as an overall measure of the β‐cell response relative to the glucose challenge (76) also showed a non-significant decrease in the patients. Besides, OGTT-based composite measures as indicators of the ability of β‐cells to compensate for insulin resistance (79–81) showed no significant reductions in the SCA2 patients relative to control individuals (Table S3).

Altogether, though all insulin sensitivity and secretion indices were reduced, and all insulin resistance indices were increased in the patients relative to control individuals, only the fasting glucose levels and QUICKI index reached nominal significance. Therefore, findings from the fasting state and the OGTT do not consistently support the assumption of altered blood glucose homeostasis in SCA2 patients with mild to moderate ataxia.

### Markers of blood glucose homeostasis show weak to moderate associations with body composition and disease severity in patients with SCA2

Given that disease severity and progression in patients with SCA2 may be influenced by metabolic factors in addition to genetic modifiers (41, 46, 59, 89), we assessed the effects of blood glucose homeostasis markers on anthropometric measures of body composition and disease severity. As expected from established associations between glucose/energy metabolism and body composition (90), fasting insulin levels, QUICKI, McAuley’s, HOMA2-%S, HOMA2-IR, and HOMA2-%β indices showed significant correlations with the body weight, waist circumference, and BMI, with more or less diffuse dispersion of data points (Table 2; Figure S1). Most of the associations between markers of glucose homeostasis and waist circumference and BMI, but none of the correlations with body weight, remained significant after correction for multiple hypothesis testing. Conversely, no significant associations were observed between the *ATXN2* CAG repeat length, disease duration, or SARA score, and the glucose homeostasis markers from the fasting state (Table S4). Comparisons across disease stages did not produce any significant results either (Figure S2). However, McAuley’s index (Mffm/I) for insulin sensitivity showed significant correlations with the age at disease onset (r= −0.239; p= 0.038), progression rate (r= 0.318; p=0.005), and INAS count (r= 0.277; p= 0.021). Besides, the TyG index for insulin resistance was significantly correlated with the progression rate (r= −0.293; p=0.008), and the HOMA2-%β index showed a weak correlation with the INAS count (r= −0.241; p=0.046) (Figure S3; Table S4). Nonetheless, only the correlation between McAuley’s index and progression rate remained significant after correction for multiple hypothesis testing (corrected p-value: 0.045).

**Table 2.** Correlation analysis between anthropometric markers of body composition and markers of blood glucose homeostasis in patients with SCA2.

|  | Body weight (kg) | Waist Circ. (cm) | BMI (kg/m<sup>2</sup>) |
| --- | --- | --- | --- |
| Triglycerides (mmol/L) | 0.207 (0.093) | 0.221 (0.073) | 0.205 (0.074) |
| FG (mmol/L) | 0.053 (0.671) | 0.163 (0.189) | 0.035 (0.764) |
| FI (μUI/mL) | 0.268 (**0.033**) | 0.334 (**0.007**) | 0.295 (**0.011**) |
| *Insulin sensitivity* |  |  |  |
| QUICKI | -0.291 (**0.020**) | -0.351 (**0.004**) | -0.342 (**0.003**) |
| Mffm/I | -0.279 (**0.026**) | -0.342 (**0.006**) | -0.313 (**0.007**) |
| HOMA2-%S | -0.215 (0.096) | -0.301 (**0.018**) | -0.283 (**0.017**) |
| *Insulin resistance* |  |  |  |
| TyG | 0.158 (0.203) | 0.222 (0.071) | 0.163 (0.157) |
| HOMA2-IR | 0.262 (**0.037**) | 0.333 (**0.007**) | 0.294 (**0.011**) |
| *Insulin secretion* |  |  |  |
| HOMA2-%β | 0.267 (**0.033**) | 0.264 (**0.035**) | 0.277 (**0.017**) |

FG– fasting serum glucose levels; FI– fasting serum insulin levels; TyG– Triglyceride-Glucose index; QUICKI– Quantitative Insulin Sensitivity Check Index; Mffm/I– McAuley index; HOMA2-%β– Homeostasis Model Assessment of Beta-Cell function; HOMA2-%S– Homeostasis Model Assessment of Insulin Sensitivity; HOMA2-IR– Homeostasis Model Assessment of Insulin Resistance; Waist Circ.– Waist circumference; BMI– body mass index. The Pearson’s correlation coefficients and the corresponding p-values are shown. Significant associations are shown in bold.

As a major modifier of clinical severity, the *ATXN2* CAG repeat length was strongly correlated with the age at onset (r= −0.713; p<0.001), SARA score (r= 0.311; p=0.006), progression rate (r= 0.746; p<0.001), and INAS count (r= 0.427; p<0.001). Besides, disease duration was associated with the SARA score (r=0.551; p<0.001). In addition, the BMI was the only anthropometric measure of body composition relevant to disease severity, showing a significant effect on the age at onset (r=0.264; p=0.024) (Table S5).

Based on these correlations, we re-tested the association between markers of clinical severity and parameters or indices of blood glucose metabolism by including the CAG repeat length, BMI, and/or disease duration as additional predictors in multiple linear regression models. Regression of the CAG repeat length, BMI, and markers of blood glucose metabolism on the age at onset produced no significant results for markers of glucose homeostasis (Table S6). Similarly, no marker of blood glucose metabolism was associated with the SARA score after correction for the CAG repeat length and disease duration (Table S7). Conversely, after correction for the CAG repeat length by multiple linear regression on the progression rate, the McAuley’s index remained significant (β(SE) = 0.016 (0.007); p=0.020), and a new association with the HOMA2-IR index emerged (β(SE) = −0.047 (0.022); p=0.038) (Table S8). No marker of blood glucose metabolism was associated with the INAS count after correction for the CAG repeat length (Table S9). None of the stronger effects obtained during multiple linear regression analysis for the markers of blood glucose metabolism remained significant after correction for multiple hypothesis testing.

Overall, association analyses indicate that blood glucose metabolism associates with body composition and has weak to moderate effects on disease severity and progression rate in patients with SCA2, with McAuley’s index of insulin sensitivity showing significant effects more consistently.

## Discussion

Given recent and future developments in experimental disease-modifying treatments for SCA2, the search for reliable and informative biomarkers with strong mechanistic correlates becomes crucially important. Despite progress made in the identification of central or peripheral imaging, electrophysiological, digital, and biochemical candidate biomarkers for SCA2 (91, 92), further efforts are needed towards the discovery and validation of new candidates. As the novel contribution of the present study, we now explored surrogate markers of blood glucose homeostasis in patients with SCA2 to identify those relevant to SCA2 pathophysiology.

Evidence supporting a pathologic role for blood glucose dyshomeostasis in polyglutamine disorders is scarce and conflicting. Reports showing altered glucose homeostasis in mouse models or patients with HD, SCA1, SCA3, SCA7 or SBMA (18, 20–24, 26, 27, 93) were challenged by additional studies showing no difference in plasma glucose levels, insulin sensitivity and secretion between HD patients and control individuals from fasting state or during the OGTT (94, 95). Similarly, we obtained no convincing evidence for an altered blood glucose homeostasis in patients with mild to moderate SCA2, apart from a subtle increase in fasting serum glucose levels along with a decreasing QUICKI index for insulin sensitivity after correction for BMI, which may be indicative of an incipient pro-insulin resistance phenotype (73, 82). These effects are reminiscent of observations in the Sca2^-/-^ mouse, where increased insulin levels in the pancreas and blood serum were described (29), as occurs in insulin resistance syndromes (96), and might reflect an Ataxin-2 partial loss-of-function due to its polyQ expansion and subsequent aggregation.

Interestingly, pancreatic β-cells, similar to neurons, are long-lived cells that show a very slow replication rate (estimated in 0.1–0.5% in the adult), remain practically quiescent throughout adulthood (97–100), and are highly sensitive to stress granule dysfunction and the aggregation of RNA‐binding proteins (101–103). However, findings from fasting state and during OGTT indicate that SCA2 patients with mild to moderate ataxia do not have any major pancreatic β-cell defect, suggesting that polyQ-expanded Ataxin-2 does not interfere substantially with insulin secretion, which might be related with the limited abundance of Ataxin-2 in pancreatic β-cells. Indeed, conventional immunohistochemistry profiling data from The Human Protein Atlas indicate that Ataxin-2 protein shows only moderate staining intensity in pancreatic endocrine cells (https://www.proteinatlas.org/ENSG00000204842-ATXN2/tissue/pancreas). Conversely, Ataxin-2 protein is highly expressed in cerebellar Purkinje cells (https://www.proteinatlas.org/ENSG00000204842-ATXN2/tissue/cerebellum#rnaseq cerebellum), which are primarily affected in patients with SCA2 (44, 104). In comparison, huntingtin (HTT) protein also shows moderate staining intensity in pancreatic endocrine cells as well (https://www.proteinatlas.org/ENSG00000197386-HTT/tissue/pancreas), but evidence indicate that polyQ expanded HTT protein aggregates in pancreatic β-cells in mouse models of HD (20, 105, 106), sequestering key insulin signaling pathway factors into the aggregates and disrupting insulin secretion (107, 108). Unfortunately, though the accumulation of polyQ-expanded Ataxin-2 protein aggregates is the pathological hallmark in the brains of patients with SCA2 (109, 110), there is no available evidence that any published study has examined aggregation of polyQ-expanded Ataxin-2 protein in pancreatic β‐cells.

Markers of blood glucose homeostasis seem to be of minor importance for body composition in patients with HD or SCA1 (22, 95), though the limited sample size in these studies may be hiding true effects. Contrary to those findings in HD or SCA1 patients, a direct correlation between HOMA-IR and BMI was reported in a larger cohort of patients with SBMA (93). Similarly, most markers of blood glucose homeostasis were significantly associated with body weight, waist circumference, and BMI in our SCA2 cohort with mild to moderate ataxia. Given that Ataxin-2 protein has been involved in lipid storage and energy metabolism (29, 60), the observed correlations suggest a conserved functional interplay between blood glucose homeostasis and body composition, to cope with the metabolic stress and systemic energy deficits posed by the continuous expression of polyQ-expanded Ataxin-2 in peripheral tissues involved in energy storage. Indeed, *Atxn2* knockout mice manifest abdominal obesity, hepatosteatosis, dyslipidemia, and altered fatty acid pathways (29, 111, 112), whereas there is a consistent decrease in cholesterol and ceramide levels in the cerebellum or spinal cord of *Atxn2*-CAG100-KnockIn mice (60, 113). In an additional study, it was shown that Ataxin-2 lentiviral overexpression in the hypothalamus is important for modulation of body weight and insulin sensitivity in *Atxn2* knockout mice (114). Besides, unintended weight loss and a progressive decrease in BMI have been identified as hallmarks of SCA2 in mouse models or human patients (41, 58, 115–118). Given the above, the described associations between markers of glucose homeostasis and body composition in our SCA2 cohort may reflect physiological compensatory efforts under conditions of metabolic stress.

Findings of brain glucose dyshomeostasis usually correlate with markers of pathology or disease severity in common neurodegenerative diseases (5, 119), and also in rare polyglutamine disorders (13, 16, 120–122). Conversely, these correlations seem to be elusive when it comes to peripheral markers of blood glucose homeostasis in polyglutamine disorders. Indeed, the CAG repeat length showed a significant correlation only with the acute insulin response in HD patients, but not with additional markers of glucose homeostasis in patients with HD, SCA1, SCA3, or SBMA (21, 22, 26, 93, 95). Besides, markers of blood glucose homeostasis appear to possess negligible significance for severity or duration of neurological manifestations in patients with HD, SCA1, or SBMA (22, 93–95), though a direct association between insulin levels and age at onset was reported in SCA3 patients (26). Likewise, we found no strong association between the CAG repeat at *ATXN2* expanded alleles, SARA score, or disease duration, and markers of blood glucose homeostasis in Cuban patients with SCA2. Besides, we failed in providing evidence for meaningful changes in glucose homeostasis across disease stages. These findings may reflect a true absence of biological effects, or may be due to a narrow range of expansions described in this study (the largest allele with 48 repeats, and those with 32 to 40 repeats representing >80% of the total distribution), or to a limited sample size. Remarkably, results from studies in animal models for HD or SCA7 suggest that only very large CAG repeat expansions can cause impaired blood glucose homeostasis and even symptoms of diabetes mellitus (20, 27, 123). Further investigations of blood glucose homeostasis in infantile/juvenile SCA2 patients with much longer CAG repeats might help to tackle this issue.

As crucial novel findings, subtle yet significant effects on the age at onset were evident for McAuley’s index of insulin sensitivity in our SCA2 cohort, whereas McAuley’s, TyG, and HOMA2-IR indices showed effects on the progression rate. In addition, McAuley’s and HOMA2-%β indices were associated with the INAS count. These results suggest that variations in blood glucose homeostasis and energy metabolism have an impact on clinical severity in patients with mild to moderate ataxia. However, the effects on the progression rate need to be verified by longitudinal studies. Notably, McAuley’s index was the blood glucose homeostasis marker more consistently associated with clinical severity. Given that McAuley’s index captures lipid-driven insulin sensitivity (85), and the confirmed involvement of altered lipid metabolism in SCA2 (60, 117), its association with markers of clinical severity might reflect effects of lipids rather than effects of insulin levels. However, no relevant associations were obtained between blood triglycerides or insulin levels and markers of clinical severity in our SCA2 cohort.

Regarding the limitations and future extension of this study, it is acknowledged that the inclusion of patients in disease stages 1 or 2 only constrains the full assessment of the effects of variations in blood glucose homeostasis on disease severity. Cross-sectional studies including presymptomatic individuals and patients in disease stage 3, along with longitudinal studies, might be very helpful for understanding the dynamic changes in blood glucose homeostasis linked to polyQ-expanded Ataxin-2 protein expression. Besides, several factors influencing glucose homeostasis, including dietary food patterns, physical activity, sleep quality, and smoking habits, among others (124, 125), were not assessed in this study. In addition, since the progression rate in SCA2 follows a non-linear trajectory (126) and should be assessed longitudinally, the ratio between the SARA score and disease duration in our study reflects the disease progression rate at presentation only. As a result, further studies should be undertaken to clarify the significance of these factors on the findings reported here. Altogether, this study indicates that SCA2 patients with mild to moderate ataxia do not have any major alterations in blood glucose homeostasis; nonetheless, it does not exclude the possibility that glucose dyshomeostasis may occur in SCA2 patients with severe ataxia or larger repeat lengths. In addition, normal variation in markers of glucose homeostasis associates with body composition and has weak to moderate modifying effects on disease severity and progression rate in patients with SCA2, with McAuley’s index showing effects more consistently. Further longitudinal studies, from pre-symptomatic to advanced disease stages, are needed to clearly describe the dynamic changes in blood glucose homeostasis across different disease stages and to assess the validity of McAuley’s index as a candidate biomarker for SCA2 clinical severity and progression.

## Acknowledgements

The authors wish to thank the patients and healthy control individuals for their cooperation with this research, and the Cuban Ministry of Public Health (MINSAP) for its vital support.

## Author contributions

Conceptualization, L.-E.A.-M., R.A.-R., D.A-.G.; methodology, R.A.-R., D.A-.G., A.A.-S., and Y.S.-R; validation, L.-E.A.-M.; formal analysis, L.-E.A.-M.; investigation, R.A.-R., D.A-G., A.A.-S., M.A.-M., Y.S.-R, D.C.-A., and A.E.-R.; resources, R.A.-R., D.A-.G., A.A.-S., and Y.S.-R; data curation, L.-E.A.-M.; writing—original draft preparation, L.E.A.-M.; writing—review and editing, R.A.-R., and D.A-.G.; visualization, L.-E.A.-M.; project administration, L.-E.A.-M., R.A.-R., D.A-.G.; funding acquisition, L.-E.A.-M. All authors have read and agreed to the published version of the manuscript.

## Competing Interests Statement

The authors declare that they have no competing interests.

## Data availability statement

The datasets used during the current study are available from the corresponding author on reasonable request.

## Supplementary

**Figure S1.**
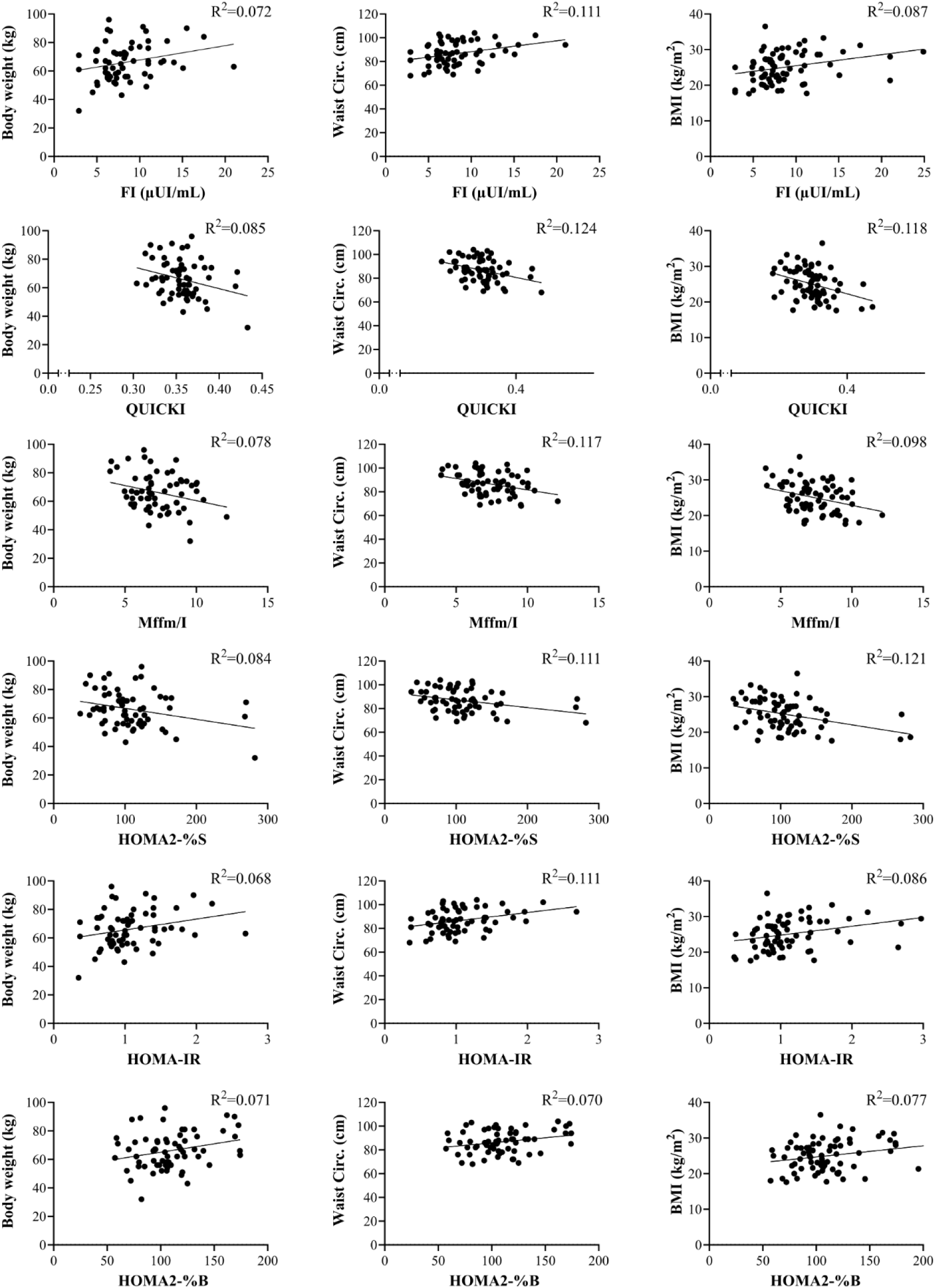
Scatter plots for the linear regression of markers of glucose homeostasis on anthropometric measures of body composition in Cuban patients with SCA2.

**Figure S2.**
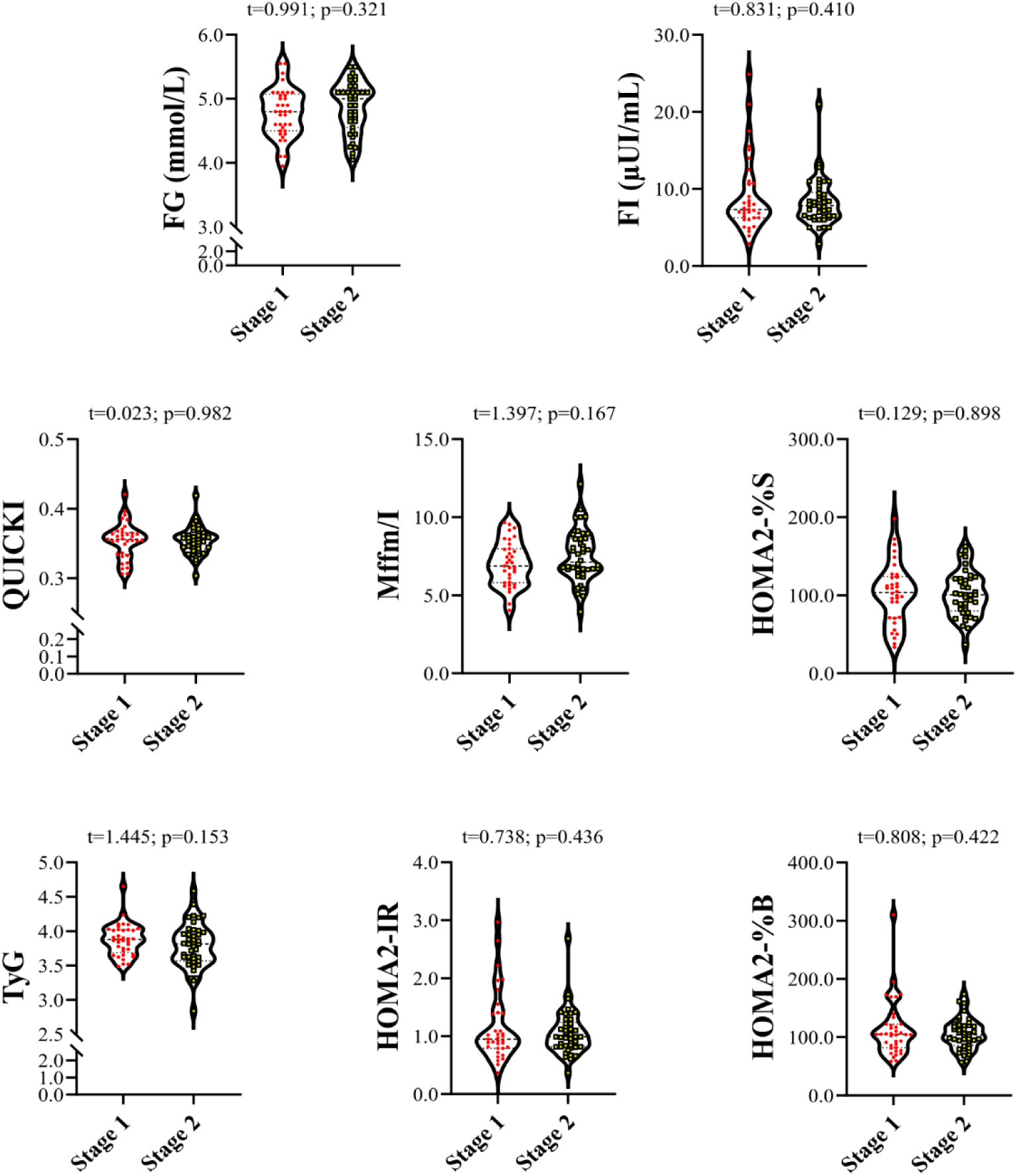
Comparison of markers of glucose homeostasis across disease stages in Cuban patients with SCA2.

**Figure S3.**
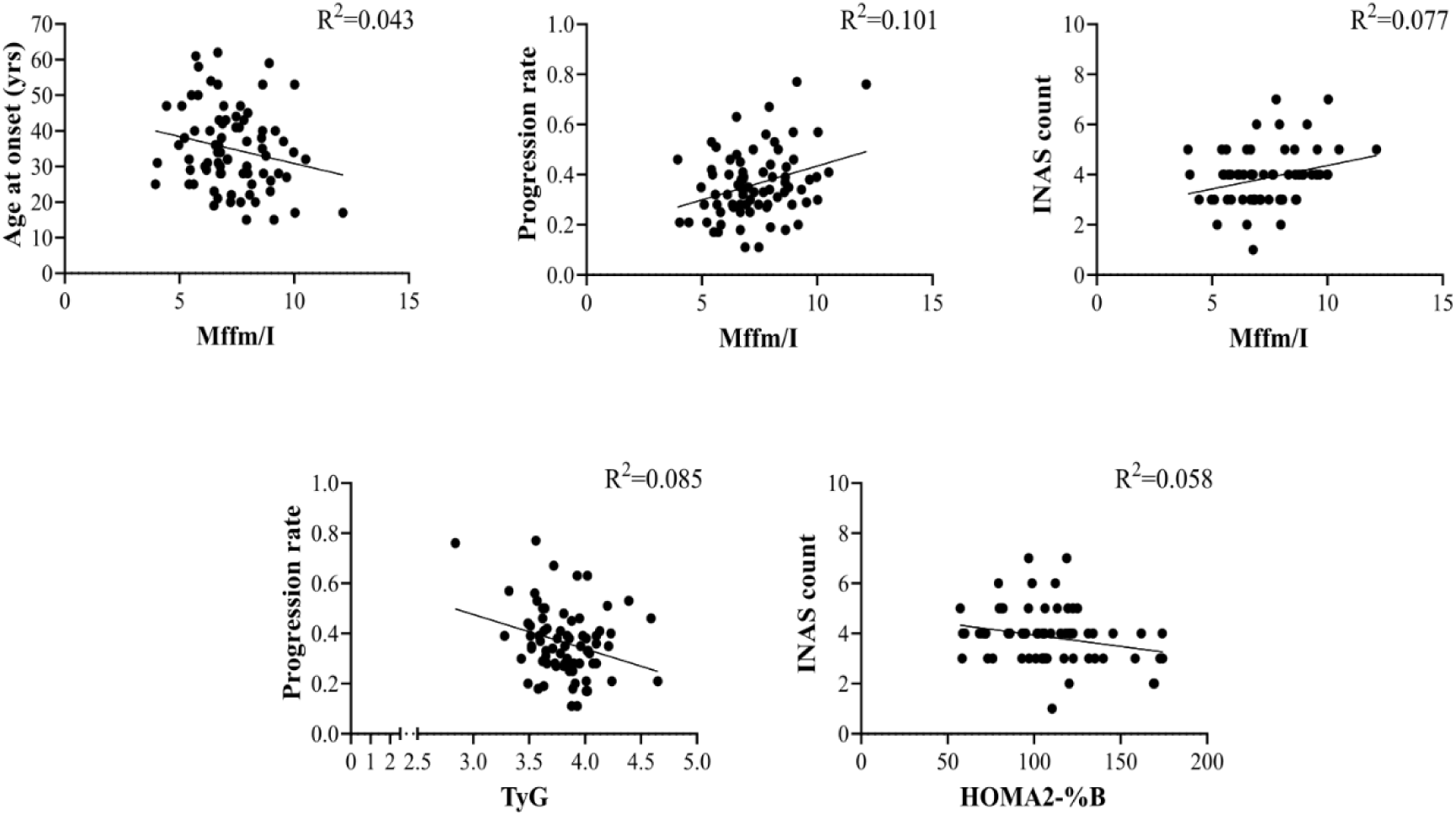
Scatter plots for the linear regression of indices of glucose homeostasis on markers of clinical severity in Cuban patients with SCA2.

**Table S1.** Comparisons between SCA2 patients and control individuals for indices of blood glucose homeostasis according to pre-defined cut-off values indicative of insulin resistance.

| | SCA2 | Controls | $\chi^2$/t-test |
| --- | --- | --- | --- |
|  | n (%) | n (%) | (p-value) |
| *Insulin sensitivity* |  |  |  |
| QUICKI (<0.35) | 23 (29.11) | 19 (22.89) | 0.816 (0.366) |
| Mffm/I ($\leq 5.0$) | 4 (5.06) | 2 (2.41) | 0.228 (0.633)<sup>a</sup> |
| HOMA2-%S (<60) | 6 (7.59) | 8 (10.96) | 0.214 (0.644) |
| *Insulin resistance* |  |  |  |
| TyG ($\geq 4.50$) | 2 (2.53) | 1 (1.20) | 0.002 (0.966)<sup>a</sup> |
| HOMA2-IR ($\geq 1.40$) | 14 (17.72) | 13 (15.66) | 0.124 (0.725) |
| *Insulin secretion* |  |  |  |
| HOMA2-%$\beta$ (<60) | 3 (3.79) | 2 (2.41) | 0.003 (0.955)<sup>a</sup> |

QUICKI– Quantitative Insulin Sensitivity Check Index; Mffm/I– McAuley index; HOMA2-%S– Homeostasis Model Assessment of Insulin Sensitivity; TyG– Triglyceride-Glucose index; HOMA2-IR– Homeostasis Model Assessment of Insulin Resistance; HOMA2-% $\beta$ – Homeostasis Model Assessment of Beta-Cell function. <sup>a</sup>– with Yate’s correction for continuity.

**Table S2.**
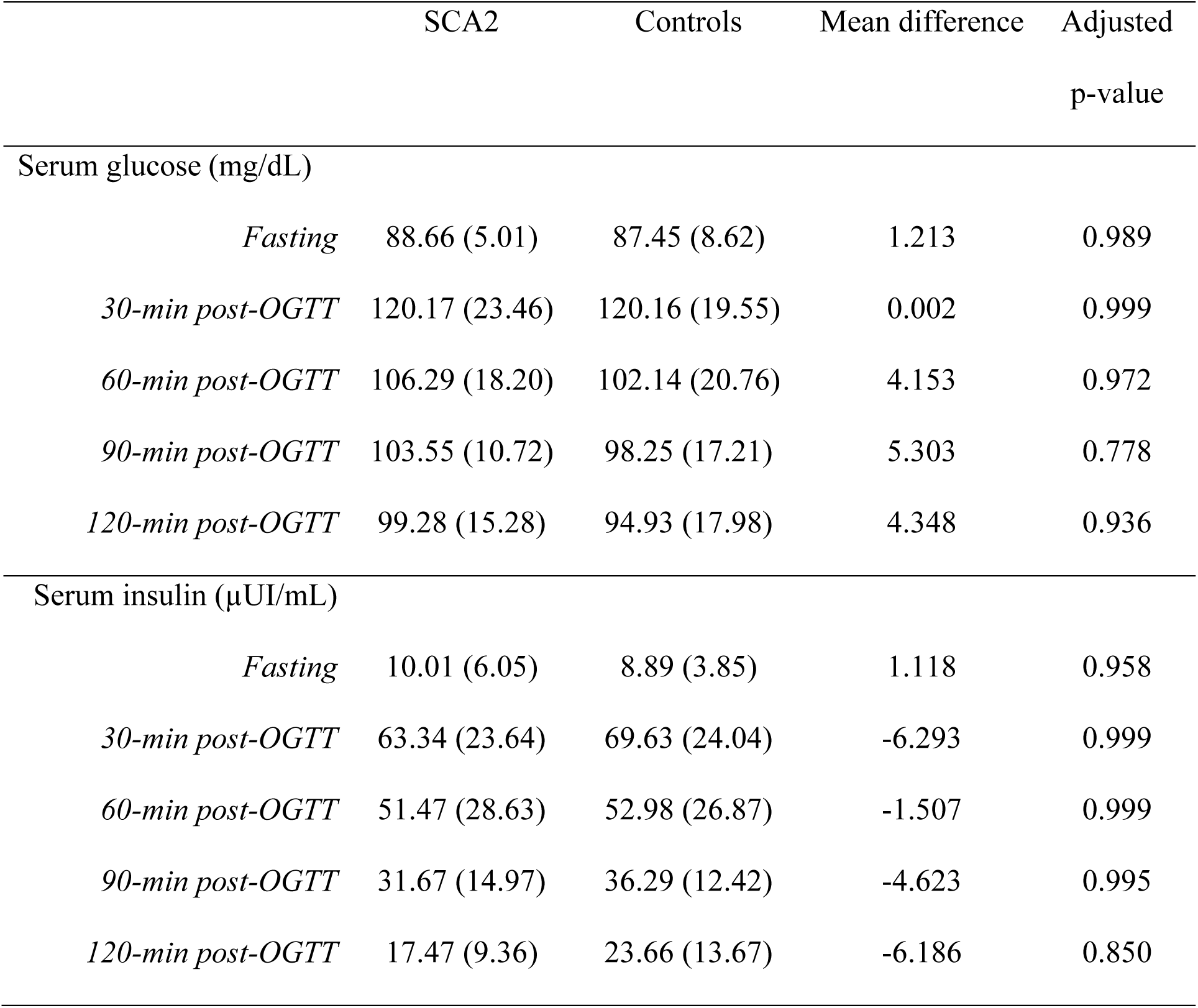
Mean comparisons between SCA2 patients and control individuals for the serum glucose and insulin levels during the oral glucose tolerance test.

|  | SCA2 | Controls | Mean difference | Adjusted<br>p-value |
| --- | --- | --- | --- | --- |
| Serum glucose (mg/dL) |  |  |  |  |
| *Fasting* | 88.66 (5.01) | 87.45 (8.62) | 1.213 | 0.989 |
| *30-min post-OGTT* | 120.17 (23.46) | 120.16 (19.55) | 0.002 | 0.999 |
| *60-min post-OGTT* | 106.29 (18.20) | 102.14 (20.76) | 4.153 | 0.972 |
| *90-min post-OGTT* | 103.55 (10.72) | 98.25 (17.21) | 5.303 | 0.778 |
| *120-min post-OGTT* | 99.28 (15.28) | 94.93 (17.98) | 4.348 | 0.936 |
| Serum insulin (μUI/mL) | | | | |
| *Fasting* | 10.01 (6.05) | 8.89 (3.85) | 1.118 | 0.958 |
| *30-min post-OGTT* | 63.34 (23.64) | 69.63 (24.04) | -6.293 | 0.999 |
| *60-min post-OGTT* | 51.47 (28.63) | 52.98 (26.87) | -1.507 | 0.999 |
| *90-min post-OGTT* | 31.67 (14.97) | 36.29 (12.42) | -4.623 | 0.995 |
| *120-min post-OGTT* | 17.47 (9.36) | 23.66 (13.67) | -6.186 | 0.850 |

**Table S3.** Descriptive statistics and mean comparisons for blood glucose metabolism indices derived from the oral glucose tolerance test, stratified by clinical status.

| Indices | SCA2 | Controls | t-test (p-value) |
| --- | --- | --- | --- |
| *Insulin sensitivity* |  |  |  |
| Gutt ISI<sub>0,120</sub> (GI) | 55.62 (33.59) | 73.18 (23.55) | 1.619 (0.119) |
| Matsuda ISI (MI) | 9.041 (4.293) | 9.888 (3.916) | 0.593 (0.558) |
| Stumvoll ISI (SI) | 0.112 (0.017) | 0.114 (0.015) | 0.328 (0.745) |
| Stumvoll MCR | 9.471 (1.432) | 9.597 (1.237) | 0.279 (0.782) |
| *Insulin secretion* |  |  |  |
| Insulinogenic index (IGI) | 0.559 (0.431) | 0.691 (0.558) | 0.685 (0.500) |
| Stumvoll PH1 | 823.20 (218.4) | 848.2 (270.5) | 0.273 (0.787) |
| Stumvoll PH2 | 222.00 (49.79) | 226.70 (57.73) | 0.239 (0.813) |
| AUC-i/g | 0.184 (0.067) | 0.247 (0.124) | 1.595 (0.131) |
| *Composite* |  |  |  |
| IGI x MI | 4.598 (2.981) | 6.735 (4.827) | 1.478 (0.152) |
| AUC-i/g x MI | 1.835 (0.702) | 2.095 (0.845) | 0.896 (0.380) |
| IGI / FI | 0.112 (0.108) | 0.115 (0.084) | 0.082 (0.936) |

MCR– metabolic clearance rate; AUC-i/g– ratio between the area under the insulin curve and the area under the glucose curve; FI– fasting insulin levels. Data are expressed as means (SD).

**Table S4.** Correlation analysis between markers disease severity and blood glucose homeostasis parameters/indices in patients with SCA2.

| Parameters/Indices | CAG | AO | SARA | DD | Progression | INAS |
| --- | --- | --- | --- | --- | --- | --- |
|  | repeat | (years) | score | (years) | rate | count |
| Triglycerides (mmol/L) | -0.014<br>(0.902) | 0.025<br>(0.824) | -0.115<br>(0.310) | 0.103<br>(0.364) | -0.149<br>(0.186) | -0.042<br>(0.726) |
| FG (mmol/L) | -0.035<br>(0.760) | 0.081<br>(0.479) | 0.025<br>(0.827) | 0.091<br>(0.420) | -0.033<br>(0.774) | 0.083<br>(0.486) |
| FI (μUI/mL) | 0.105<br>(0.371) | 0.172<br>(0.137) | -0.122<br>(0.290) | -0.121<br>(0.295) | -0.109<br>(0.345) | -0.179<br>(0.140) |
| *Insulin sensitivity* |  |  |  |  |  |  |
| QUICKI | -0.055<br>(0.643) | -0.163<br>(0.160) | 0.054<br>(0.644) | 0.037<br>(0.746) | 0.087<br>(0.453) | 0.146<br>(0.230) |
| Mffm/I | 0.188<br>(0.108) | -0.239<br>**(0.038)** | 0.191<br>(0.097) | -0.002<br>(0.989) | 0.318<br>**(0.005)** | 0.277<br>**(0.021)** |
| HOMA2-%S | -0.045<br>(0.702) | -0.134<br>(0.247) | 0.021<br>(0.856) | 0.021<br>(0.859) | 0.062<br>(0.591) | 0.156<br>(0.201) |
| *Insulin resistance* |  |  |  |  |  |  |
| TyG | -0.209<br>(0.068) | 0.170<br>(0.133) | -0.177<br>(0.117) | 0.060<br>(0.598) | -0.293<br>**(0.008)** | -0.162<br>(0.174) |
| HOMA2-IR | 0.104<br>(0.379) | 0.175<br>(0.131) | -0.115<br>(0.319) | -0.110<br>(0.343) | -0.110<br>(0.341) | -0.169<br>(0.166) |
| *Insulin secretion* |  |  |  |  |  |  |
| HOMA2-%β | 0.098 | 0.104 | -0.109 | -0.155 | -0.061 | -0.241 |
|  | (0.407) | (0.371) | (0.347) | (0.179) | (0.597) | **(0.046)** |

FG– fasting serum glucose levels; FI– fasting serum insulin levels; TyG– Triglyceride-Glucose index; QUICKI– Quantitative Insulin Sensitivity Check Index; Mffm/I– McAuley index; HOMA2-%β– Homeostasis Model Assessment of Beta-Cell function; HOMA2-%S– Homeostasis Model Assessment of Insulin Sensitivity; HOMA2-IR– Homeostasis Model Assessment of Insulin Resistance; AO– Age at disease onset; DD– Disease duration. The Pearson’s correlation coefficients and the corresponding p-values are shown. Significant associations are shown in bold.

**Table S5.** Correlation analysis between markers of body composition and markers of blood glucose homeostasis in patients with SCA2.

| Parameters/Indices | Body weight (kg) | Waist Circ. (cm) | BMI (kg/m<sup>2</sup>) |
| --- | --- | --- | --- |
| CAG repeat | -0.104 (0.415) | -0.051 (0.687) | -0.076 (0.520) |
| AO (yrs) | 0.167 (0.180) | 0.191 (0.124) | 0.264 (**0.024**) |
| SARA score | -0.221 (0.074) | -0.141 (0.257) | -0.187 (0.106) |
| DD (years) | -0.203 (0.100) | -0.239 (0.052) | -0.348 (**0.002**) |
| Progression rate | -0.210 (0.091) | -0.170 (0.173) | -0.200 (0.084) |
| INAS count | -0.205 (0.098) | -0.222 (0.074) | -0.195 (0.109) |

AO– Age at disease onset; DD– Disease duration; Waist Circ.– Waist circumference; BMI– body mass index. The Pearson’s correlation coefficients and the corresponding p-values are shown. Significant associations are shown in bold.

**Table S6.** Association of markers of blood glucose homeostasis from fasting state with the age at disease onset in multiple linear regression analysis in patients with SCA2.

| Parameters | Multiple linear model |  |  |
| --- | --- | --- | --- |
| | $R^2_{adj}$ | Regression coefficient (SE) | p-value |
| *ATXN2* expanded alleles | 0.533 | -0.033 (0.004) | **<0.001** |
| BMI (kg/m<sup>2</sup>) |  | 0.007 (0.003) | **0.013** |
| FG (mmol/L) | 0.530 | 0.020 (0.030) | 0.511 |
| FI (μUI/mL) | 0.523 | 0.005 (0.004) | 0.140 |
| *Insulin sensitivity* |  |  |  |
| QUICKI | 0.512 | -0.510 (0.602) | 0.400 |
| Mffm/I | 0.508 | -0.005 (0.008) | 0.579 |
| HOMA2-%S | 0.508 | 0.0001 (0.0001) | 0.650 |
| *Insulin resistance* |  |  |  |
| TyG | 0.527 | 0.010 (0.042) | 0.814 |
| HOMA2-IR | 0.524 | 0.044 (0.029) | 0.128 |
| *Insulin secretion* |  |  |  |
| HOMA2-%β | 0.510 | 0.0001 (0.0001) | 0.462 |

FG– fasting serum glucose levels; FI– fasting serum insulin levels; TyG– Triglyceride-Glucose index; QUICKI– Quantitative Insulin Sensitivity Check Index; Mffm/I– McAuley index; HOMA2-%β– Homeostasis Model Assessment of Beta-Cell function; HOMA2-%S– Homeostasis Model Assessment of Insulin Sensitivity; HOMA2-IR– Homeostasis Model Assessment of Insulin Resistance. The multiple linear models were based on the linear
regression of the *ATXN2* CAG repeat length and individual glucose homeostasis parameters/indices on the age at disease onset. Significant associations are shown in bold.

**Table S7.**
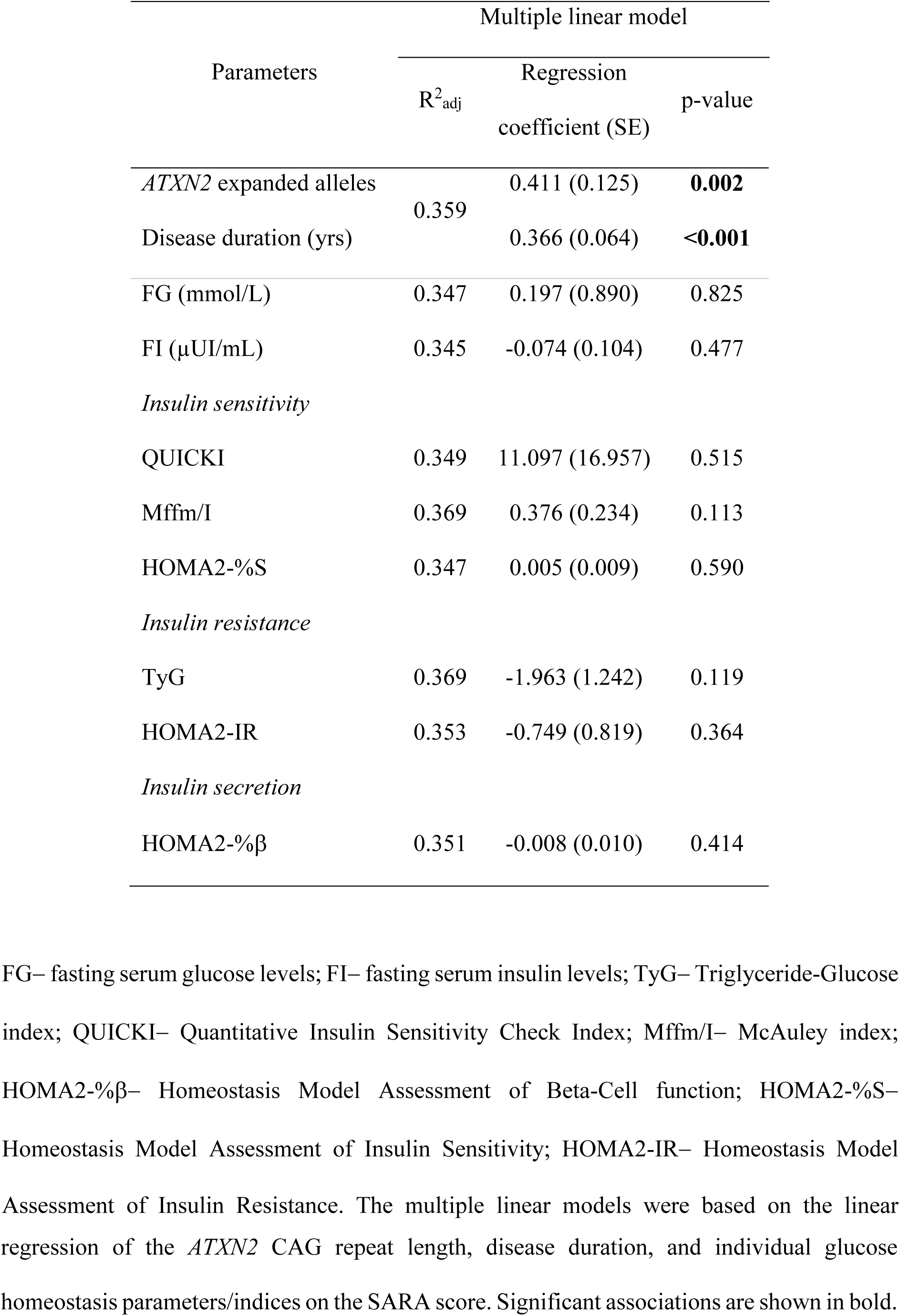
Association of markers of blood glucose homeostasis from fasting state with the SARA score in multiple linear regression analysis in patients with SCA2.

**Table S8.** Association of markers of blood glucose homeostasis from fasting state with disease progression rate in multiple linear regression analysis in patients with SCA2.

| Parameters | Multiple linear model |  |  |
| --- | --- | --- | --- |
| | $R^2_{adj}$ | Regression coefficient (SE) | p-value |
| *ATXN2* expanded alleles | 0.528 | 0.031 (0.004) | **<0.001** |
| FG (mmol/L) | 0.544 | 0.002 (0.027) | 0.949 |
| FI (μUI/mL) | 0.521 | -0.005 (0.003) | 0.069 |
| *Insulin sensitivity* |  |  |  |
| QUICKI | 0.533 | 0.659 (0.491) | 0.184 |
| Mffm/I | 0.557 | 0.016 (0.007) | **0.020** |
| HOMA2-%S | 0.528 | 0.002 (0.001) | 0.329 |
| *Insulin resistance* |  |  |  |
| TyG | 0.565 | -0.070 (0.037) | 0.062 |
| HOMA2-IR | 0.550 | -0.047 (0.022) | **0.038** |
| *Insulin secretion* |  |  |  |
| HOMA2-%$\beta$ | 0.537 | 0.001 (0.001) | 0.127 |

FG– fasting serum glucose levels; FI– fasting serum insulin levels; TyG– Triglyceride-Glucose index; QUICKI– Quantitative Insulin Sensitivity Check Index; Mffm/I– McAuley index; HOMA2-% $\beta$ – Homeostasis Model Assessment of Beta-Cell function; HOMA2-%S– Homeostasis Model Assessment of Insulin Sensitivity; HOMA2-IR– Homeostasis Model Assessment of Insulin Resistance. The multiple linear models were based on the linear
regression of the *ATXN2* CAG repeat length and individual glucose homeostasis parameters/indices on the disease progression rate. Significant associations are shown in bold.

**Table S9.** Association of markers of blood glucose homeostasis from fasting state with the INAS count in multiple linear regression analysis in patients with SCA2.

| Parameters | Multiple linear model |  |  |
| --- | --- | --- | --- |
| | $R^2_{adj}$ | Regression coefficient (SE) | p-value |
| *ATXN2* expanded alleles | 0.162 | 0.150 (0.041) | **0.001** |
| FG (mmol/L) | 0.061 | 0.597 (0.414) | 0.154 |
| FI (μUI/mL) | 0.042 | -0.053 (0.053) | 0.318 |
| *Insulin sensitivity* |  |  |  |
| QUICKI | 0.029 | 2.947 (7.778) | 0.706 |
| Mffm/I | 0.040 | 0.110 (0.117) | 0.352 |
| HOMA2-%S | 0.029 | 0.002 (0.004) | 0.694 |
| *Insulin resistance* |  |  |  |
| TyG | 0.033 | -0.195 (0.591) | 0.743 |
| HOMA2-IR | 0.042 | -0.428 (0.437) | 0.331 |
| *Insulin secretion* |  |  |  |
| HOMA2-%$\beta$ | 0.080 | -0.012 (0.006) | 0.060 |

FG– fasting serum glucose levels; FI– fasting serum insulin levels; TyG– Triglyceride-Glucose index; QUICKI– Quantitative Insulin Sensitivity Check Index; Mffm/I– McAuley index; HOMA2-% $\beta$ – Homeostasis Model Assessment of Beta-Cell function; HOMA2-%S– Homeostasis Model Assessment of Insulin Sensitivity; HOMA2-IR– Homeostasis Model Assessment of Insulin Resistance. The multiple linear models were based on the linear
regression of the *ATXN2* CAG repeat length and individual glucose homeostasis parameters/indices on the INAS count. Significant associations are shown in bold.

## Notes

### Competing Interest Statement

The authors have declared no competing interest.

## References

1. Mergenthaler P, Lindauer U, Dienel GA, Meisel A. Sugar for the brain: the role of glucose in physiological and pathological brain function. Trends Neurosci. 2013;36(10):587–97.

2. Li H, Guglielmetti C, Sei YJ, Zilberter M, Le Page LM, Shields L, et al. Neurons require glucose uptake and glycolysis in vivo. Cell Rep. 2023;42(4):112335.

3. Dienel GA. Brain Glucose Metabolism: Integration of Energetics with Function. Physiol Rev. 2019;99(1):949–1045.

4. Mainardi M, Fusco S, Grassi C. Modulation of hippocampal neural plasticity by glucose-related signaling. Neural Plast. 2015;2015:657928.

5. An Y, Varma VR, Varma S, Casanova R, Dammer E, Pletnikova O, et al. Evidence for brain glucose dysregulation in Alzheimer’s disease. Alzheimers Dement. 2018;14(3):318–29.

6. Szturm T, Beheshti I, Mahana B, Hobson DE, Goertzen A, Ko JH. Imaging Cerebral Glucose Metabolism during Dual-Task Walking in Patients with Parkinson’s disease. J Neuroimaging. 2021;31(2):356–62.

7. Arnould H, Baudouin V, Baudry A, Ribeiro LW, Ardila-Osorio H, Pietri M, et al. Loss of prion protein control of glucose metabolism promotes neurodegeneration in model of prion diseases. PLoS Pathog. 2021;17(10):e1009991.

8. Tefera TW, Steyn FJ, Ngo ST, Borges K. CNS glucose metabolism in Amyotrophic Lateral Sclerosis: a therapeutic target? Cell Biosci. 2021;11(1):14.

9. Bejanin A, Tammewar G, Marx G, Cobigo Y, Iaccarino L, Kornak J, et al. Longitudinal structural and metabolic changes in frontotemporal dementia. Neurology. 2020;95(2):e140–e54.

10. Ciarmiello A, Cannella M, Lastoria S, Simonelli M, Frati L, Rubinsztein DC, Squitieri F. Brain white-matter volume loss and glucose hypometabolism precede the clinical symptoms of Huntington’s disease. J Nucl Med. 2006;47(2):215–22.

11. Morea V, Bidollari E, Colotti G, Fiorillo A, Rosati J, De Filippis L, et al. Glucose transportation in the brain and its impairment in Huntington disease: one more shade of the energetic metabolism failure? Amino Acids. 2017;49(7):1147–57.

12. Delva A, Van Laere K, Vandenberghe W. Longitudinal Imaging of Regional Brain Volumes, SV2A, and Glucose Metabolism In Huntington’s Disease. Mov Disord. 2023;38(8):1515–26.

13. Grosch AS, Rinnenthal JL, Rönnefarth M, Lux S, Scheel M, Endres M, et al. Neurochemical Differences in Spinocerebellar Ataxia Type 14 and 1. Cerebellum. 2021;20(2):169–78.

14. Wang PS, Liu RS, Yang BH, Soong BW. Regional patterns of cerebral glucose metabolism in spinocerebellar ataxia type 2, 3 and 6: a voxel-based FDG-positron emission tomography analysis. J Neurol. 2007;254(7):838–45.

15. Ajayi A, Yu X, Wahlo-Svedin C, Tsirigotaki G, Karlström V, Ström AL. Altered p53 and NOX1 activity cause bioenergetic defects in a SCA7 polyglutamine disease model. Biochim Biophys Acta. 2015;1847(4-5):418–28.

16. Brockmann K, Reimold M, Globas C, Hauser TK, Walter U, Machulla HJ, et al. PET and MRI reveal early evidence of neurodegeneration in spinocerebellar ataxia type 17. J Nucl Med. 2012;53(7):1074–80.

17. Sone D, Sato N, Yokoyama K, Sumida K, Kanai M, Imabayashi E, et al. Striatal glucose hypometabolism in preadolescent-onset dentatorubral-pallidoluysian atrophy. J Neurol Sci. 2016;360:121–4.

18. Podolsky S, Leopold NA, Sax DS. Increased frequency of diabetes mellitus in patients with Huntington’s chorea. Lancet. 1972;1(7765):1356–8.

19. Podolsky S, Leopold NA. Abnormal glucose tolerance and arginine tolerance tests in Huntington’s disease. Gerontology. 1977;23(1):55–63.

20. Andreassen OA, Dedeoglu A, Stanojevic V, Hughes DB, Browne SE, Leech CA, et al. Huntington’s disease of the endocrine pancreas: insulin deficiency and diabetes mellitus due to impaired insulin gene expression. Neurobiol Dis. 2002;11(3):410–24.

21. Lalić NM, Marić J, Svetel M, Jotić A, Stefanova E, Lalić K, et al. Glucose homeostasis in Huntington disease: abnormalities in insulin sensitivity and early-phase insulin secretion. Arch Neurol. 2008;65(4):476–80.

22. Lalić NM, Dragasević N, Stefanova E, Jotić A, Lalić K, Milicić T, et al. Impaired insulin sensitivity and secretion in normoglycemic patients with spinocerebellar ataxia type 1. Mov Disord. 2010;25(12):1976–80.

23. Pacheco LS, da Silveira AF, Trott A, Houenou LJ, Algarve TD, Belló C, et al. Association between Machado-Joseph disease and oxidative stress biomarkers. Mutat Res Genet Toxicol Environ Mutagen. 2013;757(2):99–103.

24. Uchihara T, Takeda Y, Kobayashi T, Kasuga T, Ishikawa K, Kirei K, et al. Unexpected clinicopathological phenotype linked to small elongation of CAG repeat in SCA1 gene. J Neurol. 2006;253(3):396–8.

25. McGarry A, Hunter K, Gaughan J, Auinger P, Ferraro TN, Pradhan B, et al. An exploratory metabolomic comparison of participants with fast or absent functional progression from 2CARE, a randomized, double-blind clinical trial in Huntington’s disease. Sci Rep. 2024;14(1):1101.

26. Saute JA, da Silva AC, Muller AP, Hansel G, de Mello AS, Maeda F, et al. Serum insulin-like system alterations in patients with spinocerebellar ataxia type 3. Mov Disord. 2011;26(4):731–5.

27. Ward JM, Stoyas CA, Switonski PM, Ichou F, Fan W, Collins B, et al. Metabolic and Organelle Morphology Defects in Mice and Human Patients Define Spinocerebellar Ataxia Type 7 as a Mitochondrial Disease. Cell Rep. 2019;26(5):1189–202.e6.

28. Martin B, Golden E, Carlson OD, Pistell P, Zhou J, Kim W, et al. Exendin-4 improves glycemic control, ameliorates brain and pancreatic pathologies, and extends survival in a mouse model of Huntington’s disease. Diabetes. 2009;58(2):318–28.

29. Lastres-Becker I, Brodesser S, Lütjohann D, Azizov M, Buchmann J, Hintermann E, et al. Insulin receptor and lipid metabolism pathology in ataxin-2 knock-out mice. Hum Mol Genet. 2008;17(10):1465–81.

30. Auburger G, Gispert S, Lahut S, Omür O, Damrath E, Heck M, Başak N. 12q24 locus association with type 1 diabetes: SH2B3 or ATXN2? World J Diabetes. 2014;5(3):316–27.

31. Imbert G, Saudou F, Yvert G, Devys D, Trottier Y, Garnier JM, et al. Cloning of the gene for spinocerebellar ataxia 2 reveals a locus with high sensitivity to expanded CAG/glutamine repeats. Nat Genet. 1996;14(3):285–91.

32. Pulst SM, Nechiporuk A, Nechiporuk T, Gispert S, Chen XN, Lopes-Cendes I, et al. Moderate expansion of a normally biallelic trinucleotide repeat in spinocerebellar ataxia type 2. Nat Genet. 1996;14(3):269–76.

33. Sanpei K, Takano H, Igarashi S, Sato T, Oyake M, Sasaki H, et al. Identification of the spinocerebellar ataxia type 2 gene using a direct identification of repeat expansion and cloning technique, DIRECT. Nat Genet. 1996;14(3):277–84.

34. Elden AC, Kim HJ, Hart MP, Chen-Plotkin AS, Johnson BS, Fang X, et al. Ataxin-2 intermediate-length polyglutamine expansions are associated with increased risk for ALS. Nature. 2010;466(7310):1069–75.

35. Glass JD, Dewan R, Ding J, Gibbs JR, Dalgard C, Keagle PJ, et al. ATXN2 intermediate expansions in amyotrophic lateral sclerosis. Brain. 2022;145(8):2671–6.

36. Chiò A, Moglia C, Canosa A, Manera U, Grassano M, Vasta R, et al. Association of Copresence of Pathogenic Variants Related to Amyotrophic Lateral Sclerosis and Prognosis. Neurology. 2023;101(1):e83–e93.

37. Rubino E, Mancini C, Boschi S, Ferrero P, Ferrone M, Bianca S, et al. ATXN2 intermediate repeat expansions influence the clinical phenotype in frontotemporal dementia. Neurobiol Aging. 2019;73:231.e7-.e9.

38. Fournier C, Anquetil V, Camuzat A, Stirati-Buron S, Sazdovitch V, Molina-Porcel L, et al. Interrupted CAG expansions in ATXN2 gene expand the genetic spectrum of frontotemporal dementias. Acta Neuropathol Commun. 2018;6(1):41.

39. Kim YE, Jeon B, Farrer MJ, Scott E, Guella I, Park SS, et al. SCA2 family presenting as typical Parkinson’s disease: 34 year follow up. Parkinsonism Relat Disord. 2017;40:69–72.

40. Ross OA, Rutherford NJ, Baker M, Soto-Ortolaza AI, Carrasquillo MM, DeJesus-Hernandez M, et al. Ataxin-2 repeat-length variation and neurodegeneration. Hum Mol Genet. 2011;20(16):3207–12.

41. Almaguer-Mederos LE, Pérez-Ávila I, Aguilera-Rodríguez R, Velázquez-Garcés M, Almaguer-Gotay D, Hechavarría-Pupo R, et al. Body Mass Index Is Significantly Associated With Disease Severity in Spinocerebellar Ataxia Type 2 Patients. Mov Disord. 2021;36(6):1372–80.

42. Almaguer-Mederos LE, Aguilera Rodríguez R, González Zaldivar Y, Almaguer Gotay D, Cuello Almarales D, Laffita Mesa J, et al. Estimation of survival in spinocerebellar ataxia type 2 Cuban patients. Clin Genet. 2013;83(3):293–4.

43. Orozco Diaz G, Nodarse Fleites A, Cordovés Sagaz R, Auburger G. Autosomal dominant cerebellar ataxia: clinical analysis of 263 patients from a homogeneous population in Holguín, Cuba. Neurology. 1990;40(9):1369–75.

44. Rüb U, Schöls L, Paulson H, Auburger G, Kermer P, Jen JC, et al. Clinical features, neurogenetics and neuropathology of the polyglutamine spinocerebellar ataxias type 1, 2, 3, 6 and 7. Prog Neurobiol. 2013;104:38–66.

45. Yang L, Dong Y, Ma Y, Ni W, Wu ZY. Genetic profile and clinical characteristics of Chinese patients with spinocerebellar ataxia type 2: A multicenter experience over 10 years. Eur J Neurol. 2021;28(3):955–64.

46. Almaguer-Mederos LE, Falcón NS, Almira YR, Zaldivar YG, Almarales DC, Góngora EM, et al. Estimation of the age at onset in spinocerebellar ataxia type 2 Cuban patients by survival analysis. Clin Genet. 2010;78(2):169–74.

47. Jacobi H, du Montcel ST, Bauer P, Giunti P, Cook A, Labrum R, et al. Long-term disease progression in spinocerebellar ataxia types 1, 2, 3, and 6: a longitudinal cohort study. Lancet Neurol. 2015;14(11):1101–8.

48. Figueroa KP, Coon H, Santos N, Velazquez L, Mederos LA, Pulst SM. Genetic analysis of age at onset variation in spinocerebellar ataxia type 2. Neurol Genet. 2017;3(3):e155.

49. Almaguer-Mederos L-E, Kandi AR, Sen N-E, Canet-Pons J, Berger L-M, Key J, et al. Spinal cord phosphoproteome of a SCA2/ALS13 mouse model reveals alteration of ATXN2-N-term SH3-actin interactome and of autophagy via WNK1-MYO6-OPTN-SQSTM1. bioRxiv. 2024:2024.11.06.622233.

50. Del Castillo U, Norkett R, Lu W, Serpinskaya A, Gelfand VI. Ataxin-2 is essential for cytoskeletal dynamics and neurodevelopment in Drosophila. iScience. 2022;25(1):103536.

51. Inagaki H, Hosoda N, Tsuiji H, Hoshino SI. Direct evidence that Ataxin-2 is a translational activator mediating cytoplasmic polyadenylation. J Biol Chem. 2020;295(47):15810–25.

52. Lastres-Becker I, Nonis D, Eich F, Klinkenberg M, Gorospe M, Kötter P, et al. Mammalian ataxin-2 modulates translation control at the pre-initiation complex via PI3K/mTOR and is induced by starvation. Biochim Biophys Acta. 2016;1862(9):1558–69.

53. Liu YJ, Wang JY, Zhang XL, Jiang LL, Hu HY. Ataxin-2 sequesters Raptor into aggregates and impairs cellular mTORC1 signaling. Febs j. 2024;291(8):1795–812.

54. Nonis D, Schmidt MHH, van de Loo S, Eich F, Dikic I, Nowock J, Auburger G. Ataxin-2 associates with the endocytosis complex and affects EGF receptor trafficking. Cell Signal. 2008;20(10):1725–39.

55. Satterfield TF, Jackson SM, Pallanck LJ. A Drosophila homolog of the polyglutamine disease gene SCA2 is a dosage-sensitive regulator of actin filament formation. Genetics. 2002;162(4):1687–702.

56. Yamagishi R, Inagaki H, Suzuki J, Hosoda N, Sugiyama H, Tomita K, et al. Concerted action of ataxin-2 and PABPC1-bound mRNA poly(A) tail in the formation of stress granules. Nucleic Acids Res. 2024;52(15):9193–209.

57. Rüb U, Del Turco D, Del Tredici K, de Vos RA, Brunt ER, Reifenberger G, et al. Thalamic involvement in a spinocerebellar ataxia type 2 (SCA2) and a spinocerebellar ataxia type 3 (SCA3) patient, and its clinical relevance. Brain. 2003;126(Pt 10):2257–72.

58. Sen NE, Canet-Pons J, Halbach MV, Arsovic A, Pilatus U, Chae WH, et al. Generation of an Atxn2-CAG100 knock-in mouse reveals N-acetylaspartate production deficit due to early Nat8l dysregulation. Neurobiol Dis. 2019;132:104559.

59. Dennis AG, Almaguer-Mederos LE, Raúl RA, Roberto RL, Luis VP, Dany CA, et al. Redox Imbalance Associates with Clinical Worsening in Spinocerebellar Ataxia Type 2. Oxid Med Cell Longev. 2021;2021:9875639.

60. Canet-Pons J, Sen NE, Arsović A, Almaguer-Mederos LE, Halbach MV, Key J, et al. Atxn2-CAG100-KnockIn mouse spinal cord shows progressive TDP43 pathology associated with cholesterol biosynthesis suppression. Neurobiol Dis. 2021;152:105289.

61. Liu J, Tang TS, Tu H, Nelson O, Herndon E, Huynh DP, et al. Deranged calcium signaling and neurodegeneration in spinocerebellar ataxia type 2. J Neurosci. 2009;29(29):9148–62.

62. Halbach MV, Gispert S, Stehning T, Damrath E, Walter M, Auburger G. Atxn2 Knockout and CAG42-Knock-in Cerebellum Shows Similarly Dysregulated Expression in Calcium Homeostasis Pathway. Cerebellum. 2017;16(1):68–81.

63. Scoles DR, Meera P, Schneider MD, Paul S, Dansithong W, Figueroa KP, et al. Antisense oligonucleotide therapy for spinocerebellar ataxia type 2. Nature. 2017;544(7650):362–6.

64. Becker LA, Huang B, Bieri G, Ma R, Knowles DA, Jafar-Nejad P, et al. Therapeutic reduction of ataxin-2 extends lifespan and reduces pathology in TDP-43 mice. Nature. 2017;544(7650):367–71.

65. Zeballos CM, Moore HJ, Smith TJ, Powell JE, Ahsan NS, Zhang S, Gaj T. Mitigating a TDP-43 proteinopathy by targeting ataxin-2 using RNA-targeting CRISPR effector proteins. Nat Commun. 2023;14(1):6492.

66. 2. Diagnosis and Classification of Diabetes: Standards of Care in Diabetes-2025. Diabetes Care. 2025;48(1 Suppl 1):S27–s49.

67. Klockgether T, Lüdtke R, Kramer B, Abele M, Bürk K, Schöls L, et al. The natural history of degenerative ataxia: a retrospective study in 466 patients. Brain. 1998;121 ( Pt 4):589–600.

68. Schmitz-Hübsch T, du Montcel ST, Baliko L, Berciano J, Boesch S, Depondt C, et al. Scale for the assessment and rating of ataxia: development of a new clinical scale. Neurology. 2006;66(11):1717–20.

69. Jacobi H, Rakowicz M, Rola R, Fancellu R, Mariotti C, Charles P, et al. Inventory of Non-Ataxia Signs (INAS): validation of a new clinical assessment instrument. Cerebellum. 2013;12(3):418–28.

70. Almaguer-Mederos LE, Almaguer-Gotay D, Aguilera-Rodríguez R, González-Zaldívar Y, Cuello-Almarales D, Laffita-Mesa J, et al. Association of glutathione S-transferase omega polymorphism and spinocerebellar ataxia type 2. J Neurol Sci. 2017;372:324–8.

71. WHO. Waist circumference and waist-hip ratio: Report of a WHO expert consultation. Geneva, Switzerland; 2011.

72. Peng F, Li X, Xiao F, Zhao R, Sun Z. Circadian clock, diurnal glucose metabolic rhythm, and dawn phenomenon. Trends Neurosci. 2022;45(6):471–82.

73. Katz A, Nambi SS, Mather K, Baron AD, Follmann DA, Sullivan G, Quon MJ. Quantitative insulin sensitivity check index: a simple, accurate method for assessing insulin sensitivity in humans. J Clin Endocrinol Metab. 2000;85(7):2402–10.

74. Levy JC, Matthews DR, Hermans MP. Correct homeostasis model assessment (HOMA) evaluation uses the computer program. Diabetes Care. 1998;21(12):2191–2.

75. Gutt M, Davis CL, Spitzer SB, Llabre MM, Kumar M, Czarnecki EM, et al. Validation of the insulin sensitivity index (ISI(0,120)): comparison with other measures. Diabetes Res Clin Pract. 2000;47(3):177–84.

76. Matsuda M, DeFronzo RA. Insulin sensitivity indices obtained from oral glucose tolerance testing: comparison with the euglycemic insulin clamp. Diabetes Care. 1999;22(9):1462–70.

77. Stumvoll M, Mitrakou A, Pimenta W, Jenssen T, Yki-Järvinen H, Van Haeften T, et al. Use of the oral glucose tolerance test to assess insulin release and insulin sensitivity. Diabetes Care. 2000;23(3):295–301.

78. Seltzer HS, Allen EW, Herron AL, Jr., Brennan MT. Insulin secretion in response to glycemic stimulus: relation of delayed initial release to carbohydrate intolerance in mild diabetes mellitus. J Clin Invest. 1967;46(3):323–35.

79. Lorenzo C, Wagenknecht LE, Rewers MJ, Karter AJ, Bergman RN, Hanley AJ, Haffner SM. Disposition index, glucose effectiveness, and conversion to type 2 diabetes: the Insulin Resistance Atherosclerosis Study (IRAS). Diabetes Care. 2010;33(9):2098–103.

80. Retnakaran R, Shen S, Hanley AJ, Vuksan V, Hamilton JK, Zinman B. Hyperbolic relationship between insulin secretion and sensitivity on oral glucose tolerance test. Obesity (Silver Spring). 2008;16(8):1901–7.

81. Utzschneider KM, Prigeon RL, Faulenbach MV, Tong J, Carr DB, Boyko EJ, et al. Oral disposition index predicts the development of future diabetes above and beyond fasting and 2-h glucose levels. Diabetes Care. 2009;32(2):335–41.

82. Hrebícek J, Janout V, Malincíková J, Horáková D, Cízek L. Detection of insulin resistance by simple quantitative insulin sensitivity check index QUICKI for epidemiological assessment and prevention. J Clin Endocrinol Metab. 2002;87(1):144–7.

83. Gokcel A, Baltali M, Tarim E, Bagis T, Gumurdulu Y, Karakose H, et al. Detection of insulin resistance in Turkish adults: a hospital-based study. Diabetes Obes Metab. 2003;5(2):126–30.

84. Salazar J, Bermúdez V, Calvo M, Olivar LC, Luzardo E, Navarro C, et al. Optimal cutoff for the evaluation of insulin resistance through triglyceride-glucose index: A cross-sectional study in a Venezuelan population. F1000Res. 2017;6:1337.

85. McAuley KA, Williams SM, Mann JI, Walker RJ, Lewis-Barned NJ, Temple LA, Duncan AW. Diagnosing insulin resistance in the general population. Diabetes Care. 2001;24(3):460–4.

86. Disse E, Bastard JP, Bonnet F, Maitrepierre C, Peyrat J, Louche-Pelissier C, Laville M. A lipid-parameter-based index for estimating insulin sensitivity and identifying insulin resistance in a healthy population. Diabetes Metab. 2008;34(5):457–63.

87. Mojiminiyi OA, Abdella NA. Effect of homeostasis model assessment computational method on the definition and associations of insulin resistance. Clin Chem Lab Med. 2010;48(11):1629–34.

88. Blüher M, Malhotra A, Bader G. Beta-cell function in treatment-naïve patients with type 2 diabetes mellitus: Analyses of baseline data from 15 clinical trials. Diabetes Obes Metab. 2023;25(5):1403–7.

89. Almaguer-Mederos LE, Aguilera-Rodríguez R, Almaguer-Gotay D, Hechavarría-Barzaga K, Álvarez-Sosa A, Chapman-Rodríguez Y, et al. Testosterone Levels Are Decreased and Associated with Disease Duration in Male Spinocerebellar Ataxia Type 2 Patients. Cerebellum. 2020;19(4):597–604.

90. Liu H, Wang S, Wang J, Guo X, Song Y, Fu K, et al. Energy metabolism in health and diseases. Signal Transduct Target Ther. 2025;10(1):69.

91. Klockgether T, Grobe-Einsler M, Faber J. Biomarkers in Spinocerebellar Ataxias. Cerebellum. 2025;24(4):104.

92. Almaguer-Mederos LE, Key J, Sen NE, Canet-Pons J, Döring C, Meierhofer D, et al. Multiomics approach identifies SERPINB1 as candidate biomarker for spinocerebellar ataxia type 2. Sci Rep. 2025;15(1):42559.

93. Rosenbohm A, Hirsch S, Volk AE, Grehl T, Grosskreutz J, Hanisch F, et al. The metabolic and endocrine characteristics in spinal and bulbar muscular atrophy. J Neurol. 2018;265(5):1026–36.

94. Boesgaard TW, Nielsen TT, Josefsen K, Hansen T, Jørgensen T, Pedersen O, et al. Huntington’s disease does not appear to increase the risk of diabetes mellitus. J Neuroendocrinol. 2009;21(9):770–6.

95. Russo CV, Salvatore E, Saccà F, Tucci T, Rinaldi C, Sorrentino P, et al. Insulin sensitivity and early-phase insulin secretion in normoglycemic Huntington’s disease patients. J Huntingtons Dis. 2013;2(4):501–7.

96. Lee SH, Park SY, Choi CS. Insulin Resistance: From Mechanisms to Therapeutic Strategies. Diabetes Metab J. 2022;46(1):15–37.

97. Meier JJ, Butler AE, Saisho Y, Monchamp T, Galasso R, Bhushan A, et al. Beta-cell replication is the primary mechanism subserving the postnatal expansion of beta-cell mass in humans. Diabetes. 2008;57(6):1584–94.

98. Gregg BE, Moore PC, Demozay D, Hall BA, Li M, Husain A, et al. Formation of a human β-cell population within pancreatic islets is set early in life. J Clin Endocrinol Metab. 2012;97(9):3197–206.

99. Wang P, Fiaschi-Taesch NM, Vasavada RC, Scott DK, García-Ocaña A, Stewart AF. Diabetes mellitus--advances and challenges in human β-cell proliferation. Nat Rev Endocrinol. 2015;11(4):201–12.

100. Cnop M, Igoillo-Esteve M, Hughes SJ, Walker JN, Cnop I, Clark A. Longevity of human islet α-and β-cells. Diabetes Obes Metab. 2011;13 Suppl 1:39–46.

101. Good AL, Stoffers DA. Stress-Induced Translational Regulation Mediated by RNA Binding Proteins: Key Links to β-Cell Failure in Diabetes. Diabetes. 2020;69(4):499–507.

102. Kulkarni A, Muralidharan C, May SC, Tersey SA, Mirmira RG. Inside the β Cell: Molecular Stress Response Pathways in Diabetes Pathogenesis. Endocrinology. 2022;164(1).

103. Araki K, Araki A, Honda D, Izumoto T, Hashizume A, Hijikata Y, et al. TDP-43 regulates early-phase insulin secretion via CaV1.2-mediated exocytosis in islets. J Clin Invest. 2019;129(9):3578–93.

104. Estrada R, Galarraga J, Orozco G, Nodarse A, Auburger G. Spinocerebellar ataxia 2 (SCA2): morphometric analyses in 11 autopsies. Acta Neuropathol. 1999;97(3):306–10.

105. Sathasivam K, Hobbs C, Turmaine M, Mangiarini L, Mahal A, Bertaux F, et al. Formation of polyglutamine inclusions in non-CNS tissue. Hum Mol Genet. 1999;8(5):813–22.

106. Moffitt H, McPhail GD, Woodman B, Hobbs C, Bates GP. Formation of polyglutamine inclusions in a wide range of non-CNS tissues in the HdhQ150 knock-in mouse model of Huntington’s disease. PLoS One. 2009;4(11):e8025.

107. Li L, Sun Y, Zhang Y, Wang W, Ye C. Mutant Huntingtin Impairs Pancreatic β-cells by Recruiting IRS-2 and Disturbing the PI3K/AKT/FoxO1 Signaling Pathway in Huntington’s Disease. J Mol Neurosci. 2021;71(12):2646–58.

108. Smith R, Bacos K, Fedele V, Soulet D, Walz HA, Obermüller S, et al. Mutant huntingtin interacts with {beta}-tubulin and disrupts vesicular transport and insulin secretion. Hum Mol Genet. 2009;18(20):3942–54.

109. Seidel K, Siswanto S, Fredrich M, Bouzrou M, den Dunnen WFA, Özerden I, et al. On the distribution of intranuclear and cytoplasmic aggregates in the brainstem of patients with spinocerebellar ataxia type 2 and 3. Brain Pathol. 2017;27(3):345–55.

110. Koyano S, Uchihara T, Fujigasaki H, Nakamura A, Yagishita S, Iwabuchi K. Neuronal intranuclear inclusions in spinocerebellar ataxia type 2: triple-labeling immunofluorescent study. Neurosci Lett. 1999;273(2):117–20.

111. Kiehl TR, Nechiporuk A, Figueroa KP, Keating MT, Huynh DP, Pulst SM. Generation and characterization of Sca2 (ataxin-2) knockout mice. Biochem Biophys Res Commun. 2006;339(1):17–24.

112. Meierhofer D, Halbach M, Şen NE, Gispert S, Auburger G. Ataxin-2 (Atxn2)-Knock-Out Mice Show Branched Chain Amino Acids and Fatty Acids Pathway Alterations. Mol Cell Proteomics. 2016;15(5):1728–39.

113. Sen NE, Arsovic A, Meierhofer D, Brodesser S, Oberschmidt C, Canet-Pons J, et al. In Human and Mouse Spino-Cerebellar Tissue, Ataxin-2 Expansion Affects Ceramide-Sphingomyelin Metabolism. Int J Mol Sci. 2019;20(23).

114. Carmo-Silva S, Ferreira-Marques M, Nóbrega C, Botelho M, Costa D, Aveleira CA, et al. Ataxin-2 in the hypothalamus at the crossroads between metabolism and clock genes. J Mol Endocrinol. 2023;70(1).

115. Damrath E, Heck MV, Gispert S, Azizov M, Nowock J, Seifried C, et al. ATXN2-CAG42 sequesters PABPC1 into insolubility and induces FBXW8 in cerebellum of old ataxic knock-in mice. PLoS Genet. 2012;8(8):e1002920.

116. Dansithong W, Paul S, Figueroa KP, Rinehart MD, Wiest S, Pflieger LT, et al. Ataxin-2 regulates RGS8 translation in a new BAC-SCA2 transgenic mouse model. PLoS Genet. 2015;11(4):e1005182.

117. Rodriguez-Graña T, Rodríguez-Labrada R, Santana-Porbén S, Reynaldo-Cejas L, Medrano-Montero J, Canales-Ochoa N, et al. Weight loss is correlated with disease severity in Spinocerebellar ataxia type 2: a cross-sectional cohort study. Nutr Neurosci. 2022;25(8):1747–55.

118. Diallo A, Jacobi H, Schmitz-Hübsch T, Cook A, Labrum R, Durr A, et al. Body Mass Index Decline Is Related to Spinocerebellar Ataxia Disease Progression. Mov Disord Clin Pract. 2017;4(5):689–97.

119. Kotagal V, Albin RL, Müller ML, Koeppe RA, Frey KA, Bohnen NI. Diabetes is associated with postural instability and gait difficulty in Parkinson disease. Parkinsonism Relat Disord. 2013;19(5):522–6.

120. Herben-Dekker M, van Oostrom JC, Roos RA, Jurgens CK, Witjes-Ané MN, Kremer HP, et al. Striatal metabolism and psychomotor speed as predictors of motor onset in Huntington’s disease. J Neurol. 2014;261(7):1387–97.

121. Meles SK, Kok JG, De Jong BM, Renken RJ, de Vries JJ, Spikman JM, et al. The cerebral metabolic topography of spinocerebellar ataxia type 3. Neuroimage Clin. 2018;19:90–7.

122. Squitieri F, Orobello S, Cannella M, Martino T, Romanelli P, Giovacchini G, et al. Riluzole protects Huntington disease patients from brain glucose hypometabolism and grey matter volume loss and increases production of neurotrophins. Eur J Nucl Med Mol Imaging. 2009;36(7):1113–20.

123. Björkqvist M, Fex M, Renström E, Wierup N, Petersén A, Gil J, et al. The R6/2 transgenic mouse model of Huntington’s disease develops diabetes due to deficient beta-cell mass and exocytosis. Hum Mol Genet. 2005;14(5):565–74.

124. Mazidi M, Kengne AP, Mikhailidis DP, Toth PP, Ray KK, Banach M. Dietary food patterns and glucose/insulin homeostasis: a cross-sectional study involving 24,182 adult Americans. Lipids Health Dis. 2017;16(1):192.

125. Jarvis PRE, Cardin JL, Nisevich-Bede PM, McCarter JP. Continuous glucose monitoring in a healthy population: understanding the post-prandial glycemic response in individuals without diabetes mellitus. Metabolism. 2023;146:155640.

126. Jacobi H, Schaprian T, Schmitz-Hübsch T, Schmid M, Klockgether T. Disease progression of spinocerebellar ataxia types 1, 2, 3 and 6 before and after ataxia onset. Ann Clin Transl Neurol. 2023;10(10):1833–43.

